# Antibody responses against bacterial glycans affinity mature and diversify in germinal centers

**DOI:** 10.1101/2025.03.26.645614

**Authors:** Holly A. Fryer, Catherine Pitt, Hannah R. Frost, Nitika Kandhari, Sean Byars, Pailene S. Lim, Trang T. Nguyen, Kaneka Chheng, Natalie Caltabiano, Alana L. Whitcombe, Julianne Hamelink, Dean Andrew, Gareth Lloyd, Brian Wilson-Boyd, Nicola Slee, Jodie Ballantine, Sarju Vasani, Kathryn Girling, Liam Gubbels, Eric Levi, Karen Davies, Stuart Tangye, Jonathan Noonan, Nicole J. Moreland, Isaak Quast, Marcus J. Robinson, Stephen W. Scally, Melanie Neeland, Shivanthan Shanthikumar, Joshua Osowicki, David M. Tarlinton, Andrew C. Steer, Michelle J. Boyle, Danika L. Hill

## Abstract

Anti-carbohydrate antibodies (Abs) play crucial roles in pathogen control, but their generation remains poorly understood. By studying responses to *Streptococcus pyogenes* in humans, we reveal that the glycan-targeted response shifts from IgM towards IgG and IgA memory with age and antigen exposure across blood, spleen, and tonsils. Both natural colonization and controlled human infection with *S. pyogenes* increased class-switched B cells, with evidence of within-clone switching. Glycan-specific B cells readily participated in germinal center (GC) responses and showed robust somatic hypermutation despite a molecular signature consistent with receiving reduced T cell help. We conclude that mucosal pathogen encounters elicit glycan responses that class-switch, evolve and diversify through the GC. These findings reveal how age and infection history can influence the quality, quantity, and isotype use of glycan-specific B cells, with implications for the design and schedule of glycan-containing vaccines.

## Introduction

The surface of many pathogens is covered with polysaccharides, which are important for both their interactions with the host and as important targets of the immune response. This is exemplified by the effectiveness of some polysaccharide-containing vaccines, including for *Streptococcus pneumoniae, Salmonella* Typhi, *Neisseria meningitides,* and *Haemophilus influenza* type B^1^. However, the utility of purified polysaccharide or glycoconjugate vaccines can be constrained by inconsistent immunogenicity in infants and young children, short-lived Ab responses and hypo-responsiveness to subsequent vaccination^2–5^. Therefore, an increased understanding of the origins, location and durability of anti-glycan B cell immunity in humans is needed to improve the design and implementation of existing and novel polysaccharide-containing vaccines.

Most purified polysaccharides are thought to elicit T-independent responses in the absence of conjugation to a protein carrier. In humans, paradoxically, many glycan-specific IgM Abs have considerable somatic hypermutation, indicative of GC passage. Many anti-carbohydrate Abs that target pathogens form early in life due to similarities with epitopes on commensal microbiota^6,7^ and are therefore often referred to as ‘natural Abs’^8,9^. This response originates within gut associated lymphoid tissue (GALT) such as the Peyer’s Patch, where somatic hypermutation can occur, and the resulting B cells can then locate to regions including the splenic marginal zone^10,11^. These ‘pre-diversified’ IgM-expressing B cells can dominate the response to subsequent antigen (Ag) encounter, often in a T-independent manner^11,12^. Class-switched anti-glycan Abs are prevalent in humans but the relationship to ‘pre-diversified’ IgM cells is not well understood and whether an IgM natural Ab response becomes fixed early in life or can continually evolve in response to pathogen exposure remains unclear. Class-switching and GC formation does not require cognate T cell help in GALT^13^, a cardinal feature in other tissues. However, the rules governing B cell function at other mucosal sites such as tonsils are poorly defined yet critical for understanding protective mucosal immunity.

*Streptococcus pyogenes* is an important human-restricted pathogen and its surface is covered in Group A carbohydrate (GAC); a polysaccharide consisting of a polyrhamnose backbone with alternating GlcNAc side chains, some containing a glycerol phosphate modification^14^. GAC is conserved across all *S. pyogenes* strains and included in several glycoconjugate vaccine formulations in development^15,16^. The GlcNAc moiety within GAC is also present on other bacterial and host glycans, hence, GlcNAc is a canonical ‘natural Ab’ IgM specificity. In mice, anti-GlcNAc Abs are generated by B1 cells in response to intestinal commensals as well as *S. pyogenes* infection^16,17,18^. In humans, both anti-GAC IgG and IgM Abs abound in healthy children and adults^19,20^. We recently reported anti-GAC IgG increased after *S. pyogenes* pharyngitis in a human challenge model and maintained for ≥3 months^21^, consistent with an anamnestic response. Therefore, determining how class-switched anti-GAC Ab responses emerge, mature, are selected and diversify is highly relevant for *S. pyogenes* vaccine design.

In this study, we reveal how GAC-specific B cells across human lymphoid tissues and blood vary by tissue, age, and infection status. By directly comparing glycan and protein responses from the same pathogen, we identified both common and distinct features of the glycan- and protein-targeting responses. While a sizeable fraction of the GAC-specific B cells was unswitched and expressed markers consistent with a marginal zone origin, isotype-switched, classical memory B cell subsets became increasingly prevalent with ageing and in response to recent *S. pyogenes* exposure. Notably, tonsillar *S. pyogenes* colonization recruited both glycan and protein-specific cells into GCs to support B cell receptor (BCR) diversification. However, reminiscent of GALT immunity, glycan-specific GC cells had a distinct light zone molecular signature indicative of receiving less T cell help, which was associated with reduced ASC output relative to the protein Ag SpyCEP. Collectively, our results reveal that repeated pathogen encounter in mucosal tissues can evolve and diversify anti-glycan responses with with kinetics and programming distinct to the contemporaneous protein response. These findings suggest that glycan-containing vaccines targeting *S. pyogenes* will elicit and/or recall class-switched, somatically hypermutated memory while also highlighting the importance of considering age, infection history and glycan antigen configuration in vaccine design and vaccination schedule.

## Results

### GAC-specific memory B cells are present in the blood and spleen of adults

To investigate the B cell response to GAC in humans, we first examined blood from 14 healthy adult donors (median 47 y.o., range 23-68 y.o.), and splenic tissue from 6 adult donors (median 58.5 y.o., range 37-69 y.o.), who are all likely to have had prior exposure to the ubiquitous human pathogen *S. pyogenes*. The splenic MZ is enriched in B cells specific for polysaccharide Ags^22^, and MZ B cells dominate human B cell responses against encapsulated bacteria like *S. pneumoniae*^11^, offering potential differences in response composition to those detected in blood.

We generated fluorescent tetramers of GAC purified from bacterial cell walls^23^, and used flow cytometry to identify GAC-binding B cells. We also generated tetramers of SpyCEP, a serine protease highly conserved among global *S. pyogenes* isolates^24^, enabling direct comparison of glycan-and protein-specific B cells elicited by the same pathogen. As an additional comparator, we investigated B cell specific to haemagglutinin (HA) from the A/Michigan/2015 (H1N1) influenza virus, where B cell memory would be present from previous mucosal infection or vaccination. This approach enabled the detection of cells specific to all three Ags in all samples studied (Figure 1A,B) at frequencies above the background staining for the fluorescent Streptavidin reagents (range 1.9-265 fold above background, Supplementary Figure 1), demonstrating the universal *S. pyogene*s and influenza virus exposure in these cohorts. The frequency of Ag positive (Ag+) cells was higher for HA but equivalent between GAC and SpyCEP in both the PBMCs and spleens (Figure 1B). By sequential gating and representation analysis (Supplementary Figure 2A-D), we observed limited CD38 and CD95 expression, molecules upregulated on GC B cells and ASCs on Ag-binding cells in the spleen (Supplementary Figure 2E-F). This indicates an absence of active GC responses; therefore, these memory B cells would have likely formed from infection or vaccination at distal sites, or been formed in the spleen early in life due to microbial encounters^11,12,18^.

**Fig. 1:**
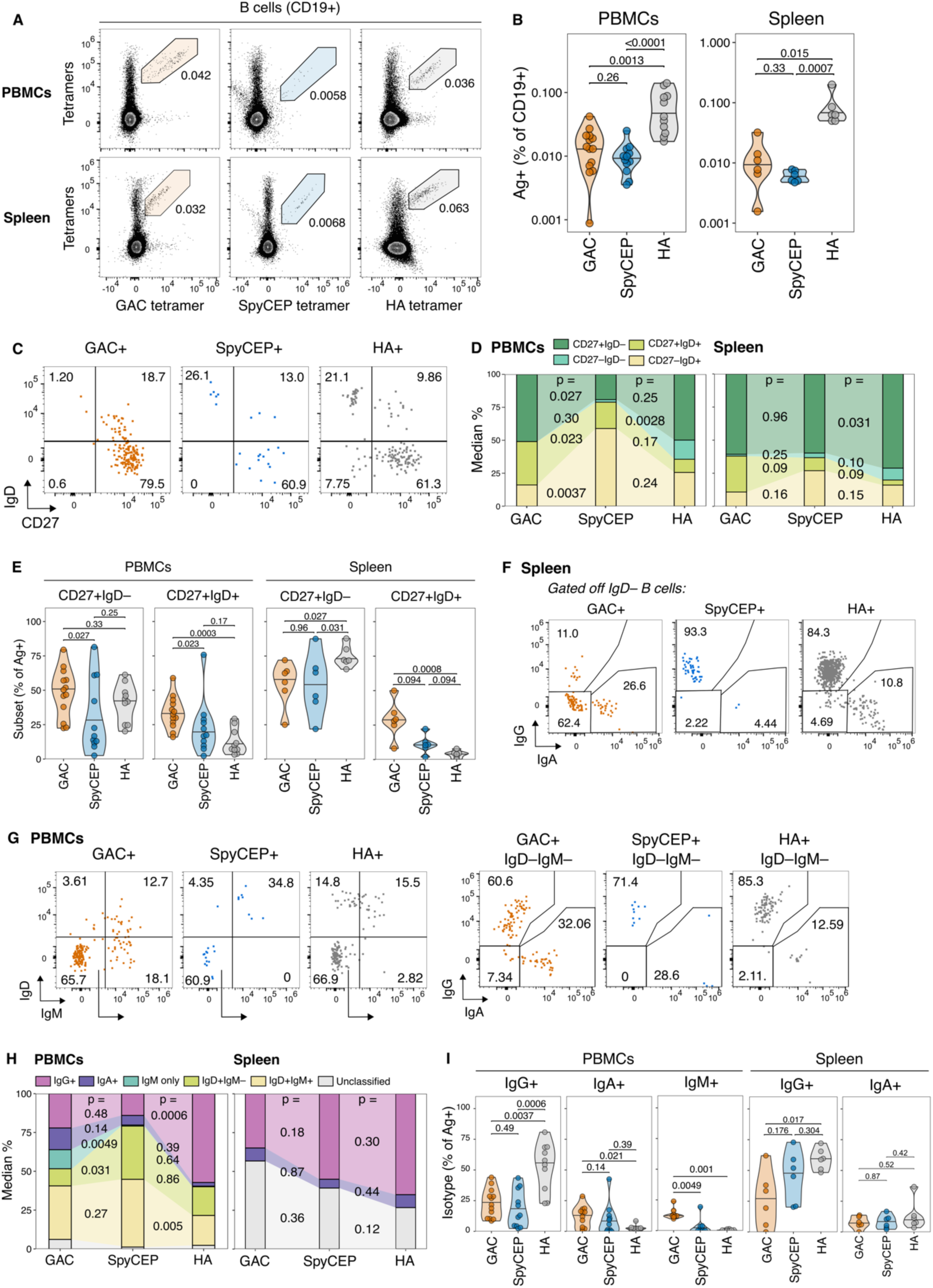
GAC-specific memory B cells are present in the blood and spleen of adults. (**A**) Representative flow cytometry plots of GAC, SpyCEP, and HA Ag dual tetramer staining on CD19+ B cells in healthy adult human PBMCs and adult splenic tissue. (**B**) Frequencies of Ag-specific (Ag+) B cells in adult PBMCs (n=14, 8♀:5♂, median 47 y.o., range 23-68 y.o., 1 unknown sex and age) and adult human spleens (n=6, 4♀:1♂, median 58.5 y.o., range 37-69 y.o., 1 unknown sex). (**C**) Representative flow cytometry plots of IgD vs CD27 staining on GAC+, SpyCEP+, and HA+ B cells, example from adult PBMCs. (**D-E**) Frequencies of GAC+, SpyCEP+, or HA+ B cell subsets in PBMCs and spleens. (**F-G**) BCR isotype distribution among GAC+, SpyCEP+, and HA+ B cells in spleens (**F**) and PBMCs (**G**). Arrows indicate hierarchical gating. (**H-I**) Frequencies of B cell subsets based on isotype within GAC+, SpyCEP+, or HA+ cells in PBMCs and spleens. Horizontal bars in violin plots indicate median values. P values were determined using two-sided non-parametric Dunn’s test with false discovery rate p-value adjustment for multiple comparisons across each population.

We co-stained with Abs against IgD and CD27 to characterize four populations among Ag-binding cells: naïve (CD27–IgD+), ‘double-negative’ (CD27–IgD–), classical memory (CD27+IgD–), and a CD27+IgD+ population that could contain IgD-only memory B cells, as well as IgD+IgM+ cells broadly consistent with a marginal zone-like phenotype in the spleen (Figure 1C). We observed some binding across all three antigens to naïve B cells, although binding of GAC to naïve B cells was the lowest of all three Ags (Supplementary Figure 2C-D). Around half of the GAC+ B cells from blood donors were of the classical memory phenotype, equivalent to their representation among HA+ B cells but significantly higher than SpyCEP+ cells (Figure 1D,E). In the spleen, most Ag+ cells had a classical memory (CD27+IgD–) phenotype (Figure 1D-E), however we also observed enrichment for unswitched CD27+IgD+ cells among GAC+ cells relative to the other Ags in both tissues (Figure 1E). We further explored isotype use of cells by staining with Abs against IgG and IgA in the splenic tissue (Figure 1F), and against IgG, IgA, IgM, and IgD in the blood (Figure 1G). Across both tissues, IgG+ and IgA+ cells were observed for all Ags, with enrichment for IgG+ among HA+ cells and IgA+ among the *S. pyogenes* Ags relative to HA (Figure 1H,I). Further analysis of non-class switched cells in the blood revealed that while IgD+IgM+ and IgD only cells were observed across all Ags; IgM-only cells were almost exclusively seen among GAC-binding cells (Figure 1H,I). In summary, while adult blood and splenic tissue contain non-class switched cells consistent with a ‘natural Ab’ response, there is a notable fraction of classical memory B cells that have isotype-switched to IgG or IgA.

### Maturation of GAC-specific B cells in the tonsil with age

Having identified that the GAC-specific B cells in the peripheral blood and spleen of adults contains class-switched memory, we next investigated the palatine tonsil, a major site of subclinical and clinical *S. pyogenes* infections (‘strep throat’)^25^. Humans typically encounter *S. pyogenes* many times throughout life such that age approximates increased cumulative pathogen exposure^26^. We used resected tonsil tissue from children (median 7 y.o., range 2-15 y.o.) and adults (median 35 y.o., range 29-41 y.o.) undergoing surgery for obstructive sleep apnoea or hyperplasia to study these responses (gating strategy Supplementary Figure 3A). All tonsil samples contained cells specific to all three Ags, indicating ubiquitous Ag exposure, with higher frequencies of GAC+ and HA+ cells in adults compared to children (Figure 2A-B).

**Fig. 2:**
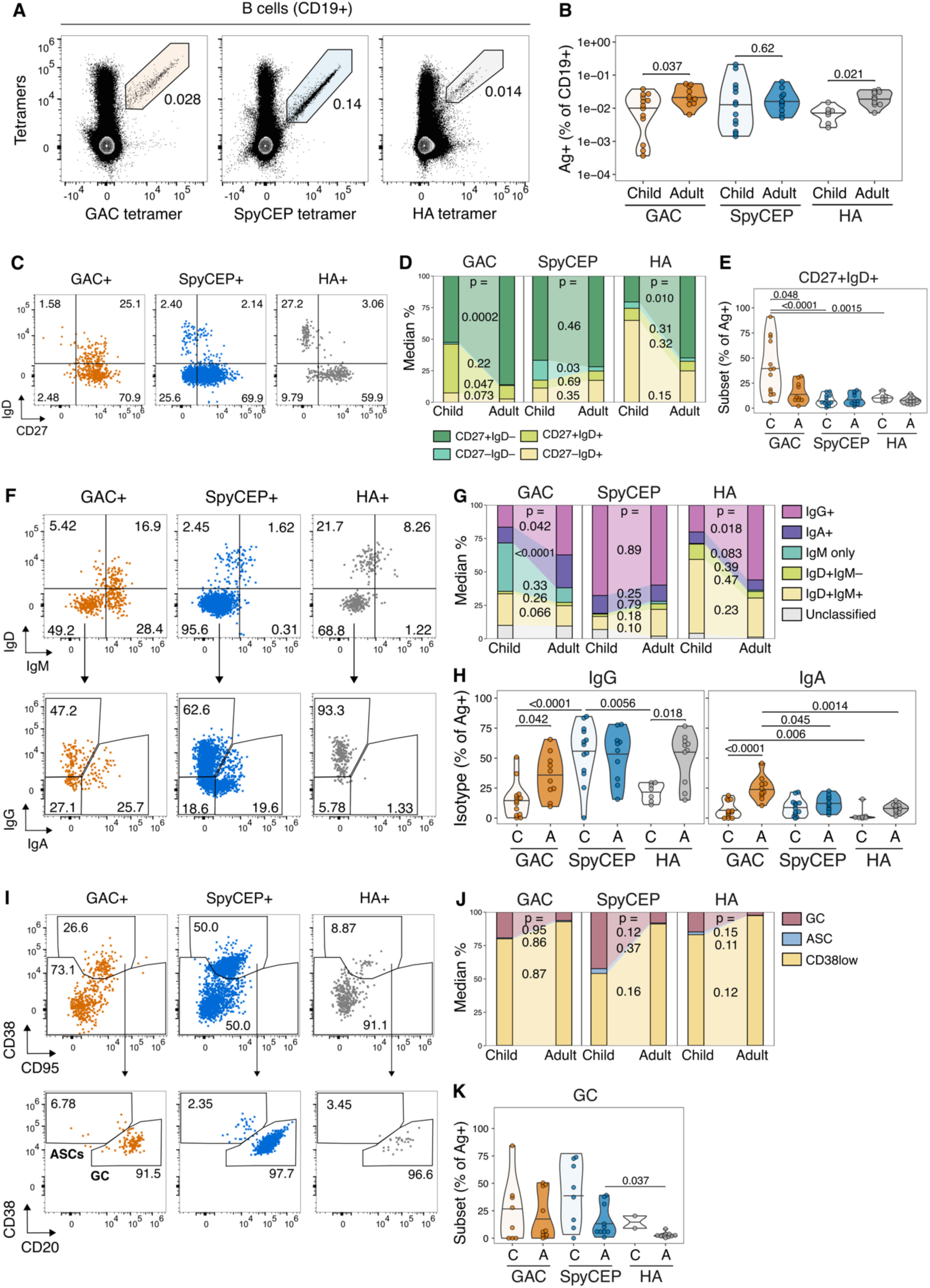
Age-related changes in GAC-specific B cells in the tonsil. (A) Representative flow cytometry plots of GAC, SpyCEP, and HA Ag dual tetramer staining on CD19+ B cells in adult human tonsils; gates correspond to Ag-specific B cells. (B) Frequencies of Ag-specific (Ag+) B cells in child (n=13, 4♀:9♂, median 7 y.o., range 2-15 y.o.) and adult (n=10, 5♀:5♂, median 35 y.o., range 29-41 y.o.) tonsils. (C) Representative flow cytometry plots of IgD vs CD27 staining on GAC+, SpyCEP+, and HA+ tonsillar B cells and associated gates. (D-E) Frequencies of GAC+, SpyCEP+, or HA+ B cell subsets in child vs adult tonsils. (F) Representative flow cytometry plots of IgD, IgM, IgG, and IgA staining on GAC+, SpyCEP+, and HA+ B cells. Arrows indicate hierarchical gating. (G-H) Frequencies of B cell subsets based on isotype within GAC+, SpyCEP+, or HA+ cells in child vs adult tonsils. (I) Representative flow cytometry plots of CD38 and CD95 staining, and CD38 vs CD20 staining indicating ASC and GC B cell gates, on GAC+, SpyCEP+, and HA+ B cells. Arrows indicate hierarchical gating. (J-K) Frequencies of GC, ASC, and CD38^low^ cells within GAC+, SpyCEP+, or HA+ B cells in child vs adult tonsils. Horizontal bars in violin plots indicate median values, P values were determined using two-sided non-parametric Dunn’s test with false discovery rate adjustment for 15 pairwise comparisons. Only p <0.05 shown in (E, H, and K).

Using CD27 and IgD to assess B cell memory composition (Figure 2C), we found broad age-related increases in classical memory (CD27+IgD–) among GAC+ and HA+ cells, but surprisingly not for SpyCEP+ cells which were abundantly represented irrespective of age (Figure 2D, Supplementary Figure 3B). In children, CD27+IgD+ cells were present in the GAC+ population but decreased in adults to a proportion similar to the other Ags (Figure 2D, E). Examining the distribution of isotypes within the Ag-specific B cells (Figure 2F), we found that proportions of IgG+ and IgA+ GAC+ but not SpyCEP+ B cells increased in adults (Figure 2G, H; Supplementary Figure 3C). Additionally, around 25% of GAC+ B cells in children were IgM+IgD–, an isotype configuration unique to the GAC+ population (Figure 2G, Supplementary Figure 3C). Class-switching to IgG and IgA also increased with age among HA+ B cells. These findings suggest that while T-cell dependent responses to bacterial protein antigens can rapidly Ab class-switch, glycan-specific B cell responses gradually evolve from predominantly IgD and/or IgM expressing cells towards isotype-switched classical memory with repeated pathogen encounters.

In contrast to the spleen, where Ag-specific GC B cells were essentially absent, GAC+ and SpyCEP+ B cells had a GC phenotype (CD38+CD95+CD20+) in some children and adult tonsils, along with a small population of ASCs expressing sufficient cell-surface BCR to bind Ag tetramers^27,28^, but did not differ between age groups (Figure 2I-K; Supplementary Figure 3D). We validated our GC and ASC gating approach by co-staining for intranuclear Ki67 and BCL6, and surface CD27 expression (Supplementary Figure 4A-D). The frequency of GAC+ and SpyCEP+ B cells with a GC phenotype varied widely (range 0-84%, Figure 2K), with many samples lacking Ag-specific GC B cells altogether, despite the presence of GCs within all tonsil samples (Supplementary Figure 4E). This variability aligns with the transient nature of GCs and suggests that GC participation requires cognate Ag availability from recent or active bacterial infection or colonization.

### *S. pyogenes* colonization increases the abundance and GC participation of glycan-specific B cells

We hypothesized that bacterial exposure was responsible for the alterations observed between children and adults in the composition of Ag-specific cells, and for the variable GC participation. To test this, we measured Ag-specific cells in an independent cohort of pediatric tonsils with and without microbiologically confirmed presence of *S. pyogenes* bacteria in the tonsil tissue. While children were not screened for *S. pyogenes* infection prior to surgery, symptomatic tonsillitis would have precluded tonsillectomy, and hence the presence of live bacteria in tonsil tissue likely reflected mild infection or asymptomatic colonization. The frequencies of GAC+ and SpyCEP+ B cells were higher in colonized compared to uncolonized tonsils, while HA+ B cells were equivalent (Figure 3A). Colonization also increased the proportion of classical memory (Figure 3B, C, Supplementary Figure 5A) and IgG+ cells (Figure 3D, E, Supplementary Figure 5B) within both GAC+ and SpyCEP+ B cells.

**Fig. 3:**
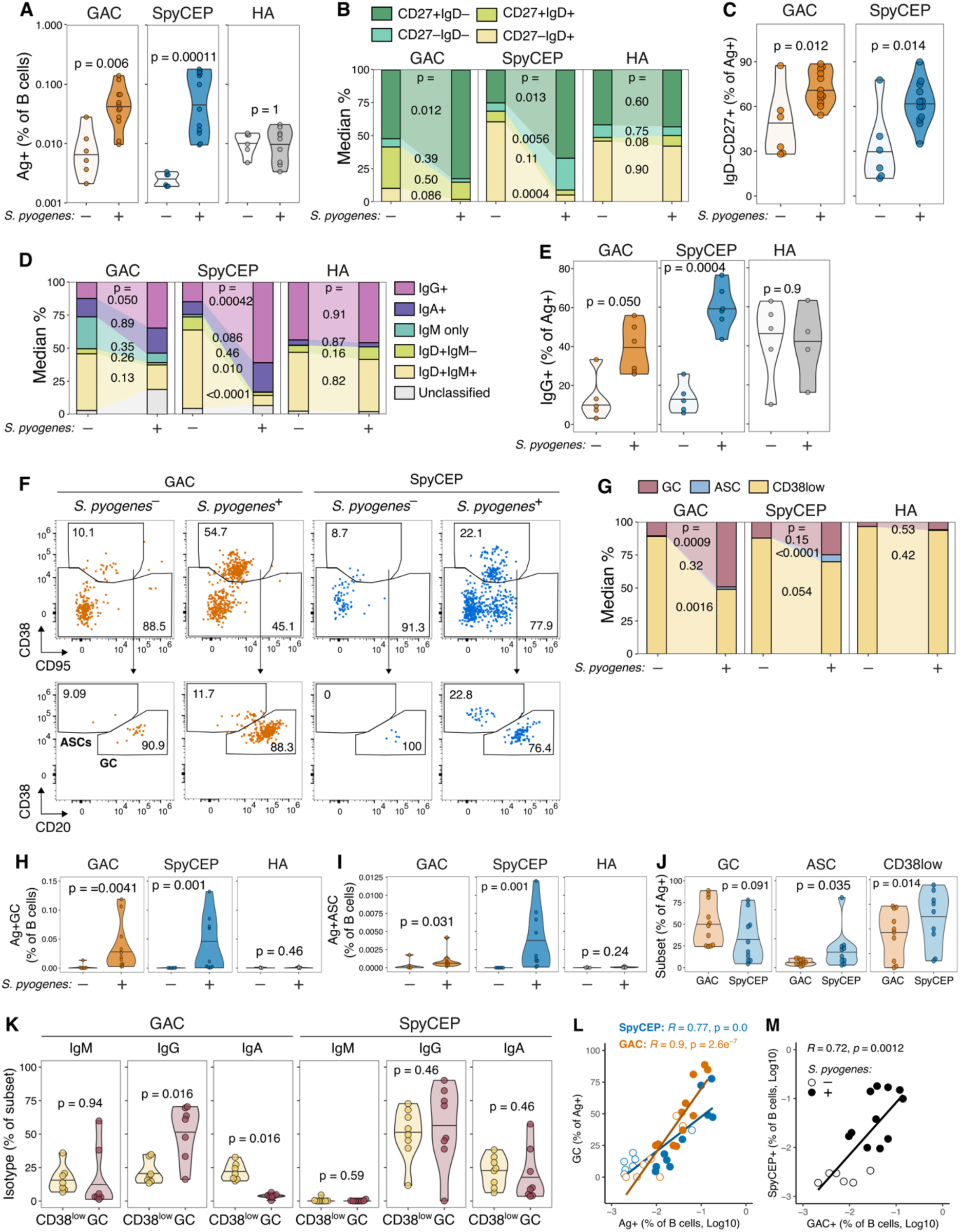
S. pyogenes colonization increases the abundance and GC participation of GAC-specific B cells in tonsils. (A) Frequencies of Ag+ B cells in pediatric human tonsils with (S. pyogenes^+^, n=12, 4♀:7♂, median 7.1 y.o, range 2.3-12.75 y.o.) and without (S. pyogenes^–^, n=7, 1♀:6♂, median 6.7 y.o, range 3.9-13.2 y.o.) colonization. (B-C) Frequencies of GAC+, SpyCEP+, or HA+ B cell subsets in tonsils with and without S. pyogenes colonization. (D-E) Frequencies of B cell subsets based on isotype within GAC+, SpyCEP+, or HA+ cells in tonsils with and without S. pyogenes colonization. (F) Representative flow cytometry plots of CD38 and CD95 staining, and CD38 vs CD20 staining ASC and GC B cell gates, on GAC+, SpyCEP+, and HA+ B cells. Arrows indicate hierarchical gating. (G) Frequencies of GC B cells, antibody-secreting cells (ASCs), and CD38^low^ cells within GAC+, SpyCEP+, or HA+ B cells. (H-I) Frequencies of GC B cells (H), or ASCs (I) (as a proportion of B cells) based on colonization status. (J) GC, ASC, and CD38low cells as a proportion of Ag+ cells in colonized tonsils (n=11), comparing GAC+ and SpyCEP+ cells. (K) IgM+, IgG+, or IgA+ cells as a proportion of CD38low or GC Ag+ cells in colonized tonsils (n=8). (A-K) Horizontal bars in violin plots indicate median values, and p values were determined using two-sided non-parametric Dunn’s test with false discovery rate adjustment for 15 pairwise comparisons. (L-M) Correlation between the frequency of GC B cells within Ag+ B cells and the frequency of Ag+ B cells for GAC and SpyCEP (L), and between GAC+ and SpyCEP+ B cells (M). Filled circles indicate S. pyogenes-colonized tonsil donors. Correlation coefficients and p values were determined using Spearman’s method (two-sided) with a linear regression line included to indicate linear trends.

Colonization increased the proportion of GAC+ B cells with a GC B cell phenotype, with a corresponding decrease in CD38^low^ B cells (Figure 3F, G, Supplementary Figure 5C). When expressed as a proportion of all B cells in the tonsil, *S. pyogenes* colonization was associated with increases in both GCs and ASCs for both GAC and SpyCEP (Figure 3H, I). We then compared the propensity of Ag-binding cells to have particular phenotypes in colonized tonsils and observed that a greater proportion of GAC+ B cells had a GC B cell phenotype compared to SpyCEP+ B cells, but more SpyCEP+ cells were enriched for ASCs relative to GAC+ cells (Figure 3J). This suggested that while GAC+ cells could readily acquire a GC phenotype, the differentiation, expansion or persistence of ASCs cells was reduced relative to SpyCEP+ cells. When comparing the surface isotype expression between GC and CD38^low^ memory populations, GAC+ GC B cells were enriched for IgG+ cells and contained fewer IgA+ cells whereas IgM was equally represented (Figure 3K). In contrast, SpyCEP+ cells showed an IgG bias and a prevalent IgA+ population, neither of which differed between the subsets (Figure 3K). For both *S. pyogenes* Ags, there was a strong correlation between Ag+ B cell frequency and the proportion of GC B cells, with both increasing proportionally in colonized tonsils (Figure 3L). In some colonized samples, more than 70% of GAC+ and SpyCEP+ B cells were of a GC phenotype, suggesting that the population expansion was closely related to GC participation. Furthermore, we observed a strong correlation between overall size of the GAC+ and the SpyCEP+ responses (Figure 3M), suggesting that protein and glycan-specific responses were elicited co-ordinately. Overall, *S. pyogenes* colonization results in BCR class switching, GC, and a larger ASC population from both glycan protein Ag targets, revealing that even mild infection or asymptomatic bacterial colonization activates and matures the B cell response to both types of antigens.

### Distinct transcriptional programs define GC responses to glycans and protein antigens

To gain further insight into B cell B cell responses in tonsils, we performed single-cell sequencing of GAC+ and SpyCEP+ IgD-B cells sorted from tonsils from four children *S. pyogenes* colonization (2.3-5.8 years old; Figure 4A, Supplementary Figure 6A). There were 7280 cells (2442 GAC+, 4838 SpyCEP+) that passed quality control, with between 935 and 3630 cells per donor. After removal of BCR and isotype genes, cells clustered into four major B cell subsets: memory, GC light zone (LZ), GC dark zone (DZ), and ASCs (Figure 4B). The distribution across major clusters was highly concordant between individuals and did not differ between Ags, except for ASCs that were infrequent among GAC+ cells (Figure 4C) consistent with earlier flow cytometry data (Figure 3J). More in-depth analysis revealed 17 distinct clusters; four clusters of memory B cells (0, 7, 8, 16), five clusters consistent with the GC light zone (LZ, 1, 2, 6, 9, 13), six clusters with the GC dark zone (DZ, 3, 5, 10, 11, 12, 15), and two ASC-related clusters (14, 4) that broadly aligned with cell type annotations from previously published tonsil single cell atlases^29^ and cell-cycle stage (Figure 4D). The transcriptional relationships between clusters were determined by analysis of selected signature genes and through hierarchical clustering of variable genes (Figure 4E-F). While one small cluster (16, ‘Bmem cycling’) of memory B cells showed moderate expression of genes associated with proliferation (e.g*. MKI67, CCNB1*), most proliferating cells were associated with the GC DZ (Figure 4E). These results suggest that proliferation within GCs is responsible for the increased frequency of Ag+ cells observed with colonization.

**Fig. 4:**
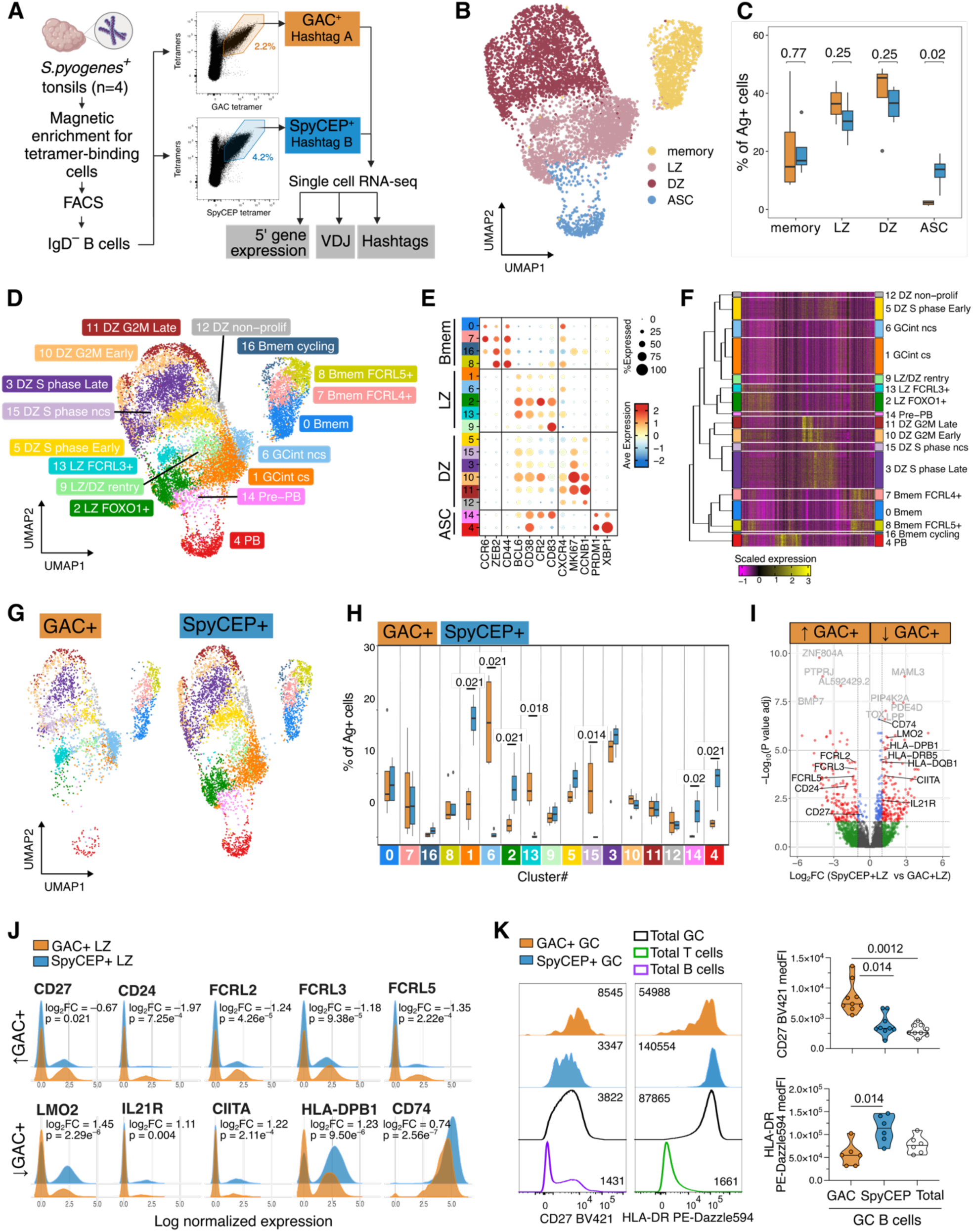
Colonization drives transcriptionally distinct responses among glycan and protein specific cells. (A) GAC+ (n=2,442 cells) and SpyCEP+ (n=4,838 cells) IgD– B cells were recovered from 4 pediatric tonsils (3♀:1♂, age range 2.3-4.69 years old) colonized with S. pyogenes, with Hashtag Abs used to barcode the Ag-specificity prior to single-cell sequencing for gene expression and BCR VDJ-seq. Figure made in BioRender. (B) Uniform Manifold Approximation and Projection (UMAP) and clustering (7,280 cells; 4 donors) into 4 major subsets, and (C) quantification of subset abundance among GAC+ or SpyCEP+ cells in each donor. Horizontal bars indicate median values, p-values determined by Dunn’s test with false discovery rate adjustment. (D) UMAP with B cell clusters and annotations shown (both Ags combined). (E) Bubble plot of selected marker genes for each major subset. Color depicts mean gene expression and dot size depicts the frequency of cells with detectable gene expression per cluster. (F) A heatmap with hierarchical clustering of the most variable genes per cluster. (G) UMAP split by Ag (2442 GAC+ cells, 4838 SpyCEP+ cells), and (H) the percentage of cells within each cluster per person per Ag. (I) Volcano plot showing differentially expressed genes between GAC+ LZ cells and SpyCEP+ LZ cells, red dots indicate genes with a log2 fold-change >1 and an adjusted p-value >0.05, (J) Selected differentially expressed genes between GAC+ and SpyCEP+ LZ cells. (K) Representative plots and quantitation of HLA-DR and CD27 median fluorescence intensity on GAC+ and SpyCEP+ GC B cells. (H, K) P-values determined by Dunn’s test with false discovery rate adjustment. Only p-values <0.05 shown. Bmem, memory B cells; GC, Germinal Center; LZ, Light zone; DZ, Dark zone; int, intermediate; PB, Plasmablast; Class-switched, cs; non-class switched, ncs.

GAC+ and SpyCEP+ B cells displayed different distributions across clusters, particularly evident in the LZ where clusters 1 (GCint class-switched (cs)) and 2 (LZ FOXO1+) were almost absent among GAC+ cells, and 6 (GCint non-class-switched (ncs)) and 13 (LZ FCRL3+) that were increased instead (Figure 4G, H). For T-dependent Ags, the LZ is where B cells present cognate Ag via MHC-II to receive help via CD40 co-stimulation and cytokines such as IL-21^30–33^. To explore if there were unique aspects to glycan-specific GC B cells, we determined the differentially expressed genes between GAC+ and SpyCEP+ cells in the LZ (clusters 1, 2, 6, 9, 13 combined, Figure 4I, Supplementary Data file 1). Among the 164 genes that were more highly expressed on GAC+ cells were *CD24* and *CD27*, molecules that can positively regulate B cell activation^34^. In contrast, GAC+ cells also expressed more *FCRL2*, *FCRL3* and *FCRL4* (Figure 4I, J), genes commonly associated with inhibition of B cell activation^35^ that shown to be upregulated on unswitched GC B cells^29^ (Figure 4I,J). Among the 173 genes lowly expressed on GAC+ LZ cells relative to SpyCEP+ were the LZ-associated genes *LMO2* and *IL21R*^29,36^*, as well as* lower expression of genes associated with Ag-presentation (e.g. *HLA-DPB1*, *CD74*, *CIITA*). These transcriptional differences were maintained in DZ cells and were particularly different between cluster 2 (LZ FOXO1+) and 13 (LZ FCRL3+) (Supplementary Figure 6B-D, Supplementary Data file 2). We confirmed the increased expression of CD27 and decreased expression of HLA-DR on GAC+ GC B cells by flow cytometry (Figure 4K). Together, this transcriptional signature reveals that while GAC+ cells can participate in the GC, they do so in a way consistent with having diminished capacity to solicit or respond to T cell help.

### GCs drive diversification of the glycan-specific B cell response

Having identified a unique molecular signature of GAC+ GC B cells, we next investigated whether isotype switching and somatic hypermutation among cells for which a single paired heavy and light chain B cell receptor (BCR) sequence was recovered (GAC+ 1381 cells, SpyCEP+ 2240 cells). Consistent with the young age of donors from whom the cells were isolated from, GAC+ cells were strongly enriched for B cell receptors expressing *IGHM*. Among *IGHG* expressing GAC+ cells there was a bias towards *IGHG1* (Figure 5A), rather the T-independent polysaccharide response-associated *IGHG2*^37,38^. In contrast, almost all SpyCEP+ cells expressed *IGHG1* (Figure 5A). This isotype bias was not responsible for the UMAP clustering, as consistent clustering patterns were observed across all isotypes (Figure 5B). Among GAC+ cells, we observed equal representation of isotypes among memory, GC and ASC populations, except for IGHA1, which was rare among GC cells but enriched among ASCs (Figure 5C).

**Fig. 5:**
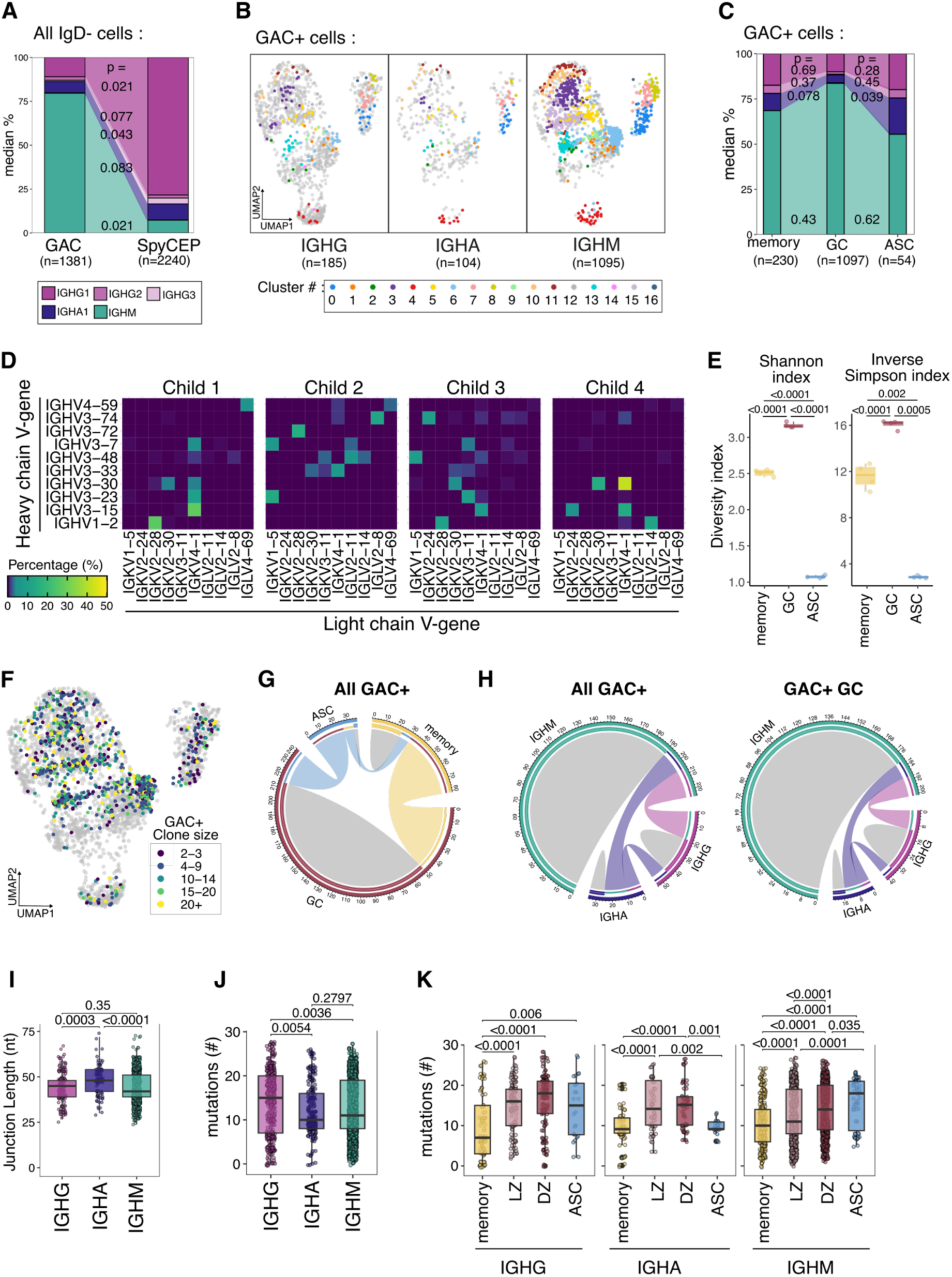
GAC-specific B cells undergo clonal expansion and somatic hypermutation in the tonsil. (A) Median proportions of cells expressing each isotype separated by Ag specificity across 4 donors. (B) UMAP showing where GAC+ B cells of each isotype map across the 17 clusters, shown against all cells (grey dots n=7280 cells). Number of cells indicated below. (C) Median proportions of GAC+ cells expressing of each isotype separated by cell type across 4 donors. (D) Heat map depicting heavy and light chain V-gene usage of GAC-specific B cells across the 4 pediatric tonsil donors. V-genes present in more than 5% of cells in any donor shown. (E) Shannon index and inverse Simpson index compared across cell types. (F) UMAP of total cells (grey, 7280 cells) with clonally expanded (≥2 cells/clone, 1043 cells) GAC+ cells highlighted. (G) Chord diagrams of clonal sharing among expanded clones (≥2 cells/clone), split by phenotype and isotype for all GAC+ cells (264 unique clones) or isotype among GAC+ cells (243 unique clones) within GC B cells. Numbering indicates the number of clones per category, grey and colored arcs represent clones with identical or shared fates, respectively. (H) Junction length in nucleotides split by isotype. (I) The combined number of heavy chain mutations in CDR1 and CDR2 split by isotype. (J) The combined number of heavy chain mutations in CDR1 and CDR2 split by isotype and compared across cell subsets. P-values determined using Dunn’s test with false discovery rate adjustment. ASC, antibody secreting cells; GC, germinal center; LZ, Light Zone; DZ, Dark Zone; CDR, complementarity-determining region.

V-gene biases and public clonotypes can be hallmarks of other glycan-specific B cells in humans and mice^11^, however we observed no public clones and a diverse repertoire of heavy and light chain use among GAC+ cells within and between individuals, with only *IGVK4-1* showing modest enrichment across all 4 donors (Figure 5D). When BCR clonal diversity was compared across cell subsets, the GC population showed greater diversity than memory and ASC populations, indicating that many different GAC+ B cell clones are recruited to the GC population after Ag encounter (Figure 5E). Polyclonality was a common finding in analyzed individuals, but notable clonal expansions did occur across memory, GC, and ASC populations (n= 264 total clones, Figure 5F). As such, while GC contained many unique expanded clones, others were shared across all three population; approximately 60% of expanded memory B cell clones and 70% of expanded ASC clones were shared with GC (Figure 5G). Both among all GAC+ cells, and among those exclusively within GC B cells, clonal sharing was present between isotypes, with 38% of IgG and 42% of IgA clones shared with IgM (Figure 5H). Consistent with the dominance of IgM+ cells in these individuals, most expanded clones were IgM-only. (Figure 5H). Also, most GAC+ cells had heavy chain junction lengths of around 50 nucleotides— a difference to the comparatively short junction lengths reported for other polysaccharide Abs^39,40^— and these were longer for IGHA compared to IGHG or IGHM (Figure 5I). The clonal relationships we observed indicate a shared origin for IgG and IgM responses with class switching occurring infrequently. This suggests that multiple *S. pyogenes* exposures are likely required to achieve the IgG-biased response observed in adult tonsils.

Mutation is another cardinal feature of response evolution. We assessed SHM in complementarity-determining regions (CDR) 1 and 2 and observed that IgG sequences were more mutated than their IgA or IgM counterparts (Figure 5J). Across all isotypes, memory B cells had fewer mutations than GC compartmentalized cells (Figure 5K). Within GC B cells, there was a trend towards higher mutations in DZ compared to LZ, with the strongest effect seen among IgM cells (Figure 5K). The ASC population was enriched for highly mutated clones relative to the GC only among IgM cells, with IgA ASCs showing equivalent numbers of mutations to memory (Figure 5K). Together, these data indicate that while the B cell response to a *S. pyogenes* glycan is diverse between individuals, common features include higher SHM loads within GC and ASC populations, irrespective of isotype, suggestive of affinity-based selection.

### Circulating classical memory B cells expand following controlled human infection

The proportions of *S. pyogenes*-specific B cells and ASCs increased within the tonsil in colonized individuals, however, it was unclear whether these differences reflected the influence of a single infection or a state of chronic carriage. To assess the short-term influence of infection, we examined serum Abs and peripheral blood B cell populations in adults from an *S. pyogenes* human challenge model (Figure 6A)^41^. In this cohort, we showed previously that some trial participants who developed symptomatic pharyngitis exhibited increases in serum IgG titres to both GAC and SpyCEP after experimental infection^21^. We selected 10 participants who retained increased (fold change >1.25) serum anti-GAC IgG at 3 months post-challenge (Figure 6B), many of whom also responded to SpyCEP (Figure 6C). These antibody titres remained stable between 1 and 3 months, suggesting that infection induced a sustained Ab response. We identified B cells specific for GAC, SpyCEP, and HA in PBMCs from all 10 participants (Figure 6D, E). GAC+ and SpyCEP+ cells increased after the challenge, while HA+ cells did not. Although the single infection did not drive significant phenotype or isotype changes (Figure 6F, G, Supplementary Figure 7A, B), IgM+ cell representation within GAC+ B cells decreased at 3 months relative to pre-challenge (Figure 6H). The increase in B cells binding the *S. pyogenes* Ags correlated with increased Ag-specific serum IgG to both Ags at 3 months relative to pre-challenge (Figure 6I). Thus, while the magnitude of Ab and memory B cell responses to both GAC and SpyCEP increased after a single infection, the observed maturation and diversification of this response between children and adults likely reflects the cumulative result of numerous pathogen encounters and GC responses throughout life.

**Fig. 6:**
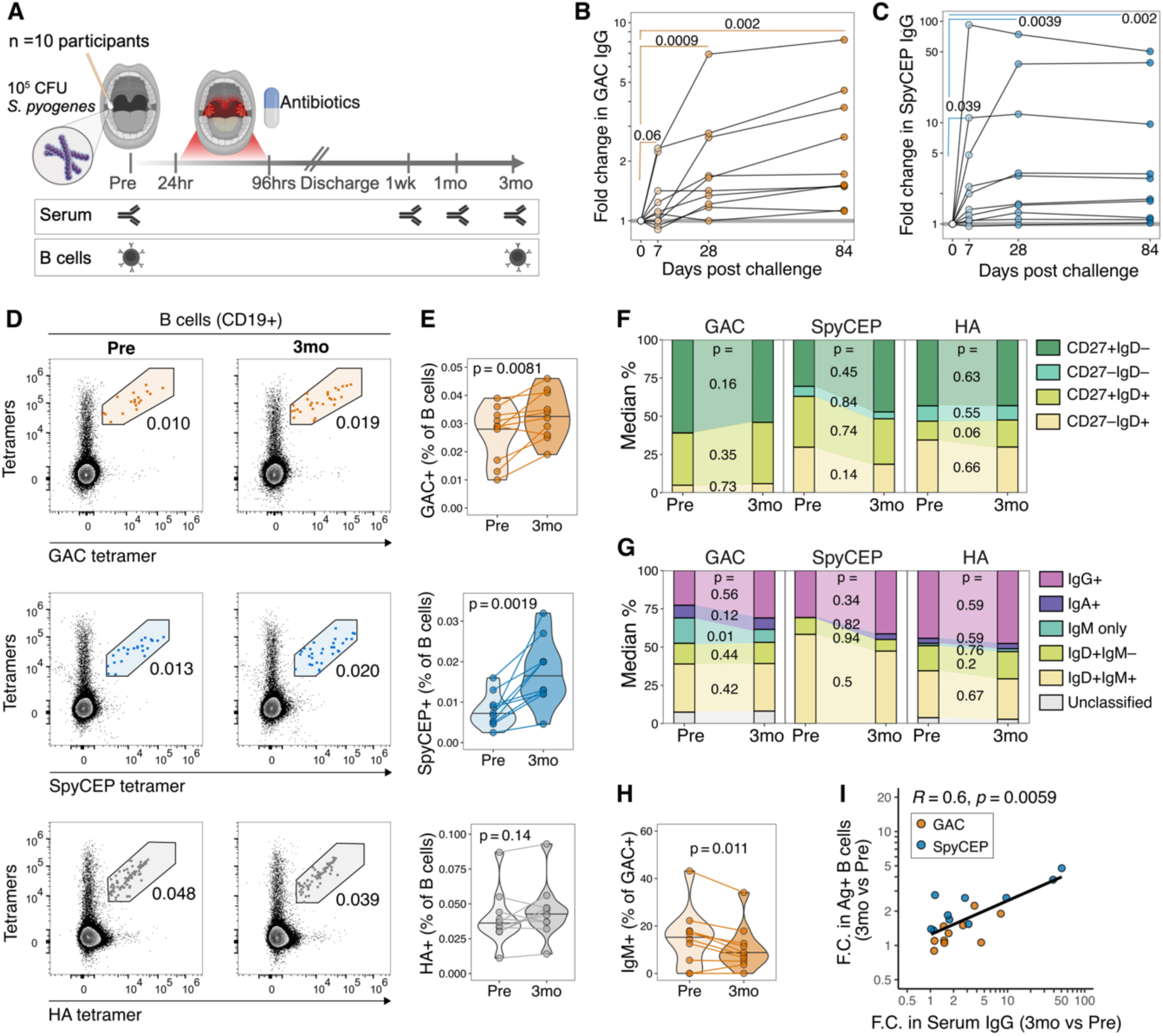
Circulating S. pyogenes glycan and protein Ag-specific antibodies and memory B cells expand following controlled human infection. (A) Schematic overview of controlled human infection study timeline and serum and PBMC (B cell) samples analyzed in this study (n=10 participants, 6♀:4♂, median 27.5 range 20.9-38.6 years old), made in Biorender. (B-C) Serum IgG responses for (B) GAC and (C) SpyCEP at pre-challenge (0 days), 1week (discharge+ 7 days), 1month (discharge+ 28 days), and 3months (discharge+ 84 days) post infectious challenge. (D) Representative flow cytometry plots of GAC, SpyCEP, and HA Ag dual tetramer staining on CD19+ B cells. (E) Frequencies of GAC+, SpyCEP+, and HA+ B cells pre- and 3 months post-challenge. (F-G) Median proportions of (F) cell subsets or (G) isotypes within GAC+, SpyCEP+, or HA+ B cells, comparing pre- and 3mo post-challenge. (H) Frequencies of IgM+ cells within GAC+ B cells, pre- and 3 months post-challenge. (I) Correlation between the fold change in Ag+ B cells and the fold change in serum Ag+ IgG at 3mo post-vs pre-challenge, for SpyCEP and GAC. Color indicates Ag specificity. Correlation coefficient and p value were determined using Spearman’s method with a linear regression line included to indicate linear trends. Horizontal bars in violin plots indicate median values. P values were determined using two-sided Wilcoxon signed-rank test for paired two-group comparisons.

We conclude that infection elicits robust immune responses against both protein and glycan antigens, with glycan-specific responses displaying a distinct and reduced reliance on cognate T cell help. Furthermore, bacterial exposure enhances the quantity, class switching, and diversity of glycan-specific B cells through a GC-dependent pathway.

## Discussion

In non-mucosal tissues like the spleen and lymph nodes, GC participation requires T cell help, with IgM-expressing cells usually receding early after response initiation, often preceding GC recruitment^42^. If T-independent GC responses form in such tissues, they are typically small and transient, with exiting B cells carrying few somatic mutations^43–45^. In contrast, GALTs such as Peyer’s Patches break many of these rules, including the need for cognate peptide presentation and CD40 signaling from T cells^46–48^. This can lead to Ag-independent BCR pre-diversification and the selection of higher-affinity clones^12,49^, as seen in T-independent responses to commensal gut microbes^12,50^ and hypermutated responses to *S. pneumoniae* capsular polysaccharides^11^ and *Klebsiella pneumoniae* O-antigen^51^. While our findings do not exclude a role for BCR pre-diversification in the B cell response to the *S. pyogenes* glycan GAC, we do show that infection recruits glycan-specific B cells into active GCs as efficiently as protein-specific cells. In the resected tonsils, the absence of clinically visible bacterial infection coupled with the high mutational load observed in GC B cells suggest that these responses do not follow the short-lived GC trajectory typical of T-independent responses. Together with adult PBMC analyses, our data support a paradigm wherein the GC is the major locale in which glycan-specific responses evolve. Beyond revealing fundamental principles of human anti-glycan responses, such knowledge is crucial in facilitating the ongoing efforts to develop *S. pyogenes*-specific vaccines aiming to optimally harness glycan-specific immunity as a means to control disease.

Glycan-specific B cells could engage in GC responses in colonized tonsils through several mechanisms. Firstly, bacterial glycans such as GAC are anchored to the cell membrane and B cells are able to engulf whole bacteria through receptor-mediated endocytosis^52^. This would enable glycan-specific cells to process and present peptides from other Ags and receive cognate T cell help. In support of this, we observed a strong correlation between glycan- and protein-specific GC populations, suggesting these responses are intertwined; we and others have already shown correlations between GAC and SpyCEP serum IgG^53,54^. The IgG-expressing B cells we detected here were almost exclusively IgG1, a hallmark of a T-dependent glycan response^55,56^. At the same time, GAC responses showed clearly distinct kinetics and our data support the notion that GAC-specific B cells receive less T cell help, indicated by low amounts of molecules and genes associated with T cell help (e.g., IL-21R) and T:B interaction (e.g., MHCII)^33,42^. Consequently, despite readily participating in GCs, it would seem GAC responses have a degree of T cell-independence. Furthermore, class switching from IgM to IgG was infrequent among GC B cell clones, suggesting limited exposure to T cell-derived class-switching cytokines, such as IL-21, prior to GC entry. Mechanistically, the combination of GAC Ag density and the repeating sugar units in GAC may favour strong BCR cross-linking, a factor that can promote GC B cell differentiation without T cell help^43^. Furthermore, *S. pyogenes* lipoteichoic acid is a ligand for TLR2, expressed on B cells, and TLR signaling can synergise with BCR signaling to sustain GC responses in the absence of T cells^57^. These insights into the GC diversification and selection are relevant to acute rheumatic fever, a rare but serious autoimmune sequela of *S. pyogenes* infection, where GAC has been implicated as a molecular mimicry trigger^58,59^. Patients with acute rheumatic fever exhibit higher anti-GAC Ab titers than healthy individuals ^19,53^, with some anti-GAC Abs displaying cross-reactivity with GlcNAc-containing heart valve glycoprotein^58,59^. Given that GCs are the source of auto-Abs in other autoimmune diseases^60^, this suggests GC involvement in development of pathogenic anti-GAC responses. The underlying phenotypic and transcriptional signatures we have identified provide further support for the concept that T cell help drives anti-glycan immunity in response to bacteria.

Polysaccharide-containing vaccines are important public health tools for preventing *Haemophilus influenzae* type B, pneumococcal and meningococcal disease^56^. However, age- and exposure-related changes in B cell responses dramatically impact vaccine immunogenicity; for example, there is a window in early life when pneumococcal conjugate vaccines can elicit a T-dependent Ab response, but this closes once a non-protective IgM+ marginal zone (MZ) response becomes established^61^. These glycan-specific MZ B cells are thought to capture available Ag and rapidly differentiate into short-lived plasmablasts, thereby inhibiting *de novo* responses from follicular B cells^61^. Our data support the existence of an *S. pyogenes* glycan-specific CD27+IgD+IgM+ population that is phenotypically consistent with splenic marginal zone B cells^22^ and was also present in peripheral blood and tonsil tissue. This population proportionally declined with age and colonization, coinciding with an increase in class-switched IgG B cells that underwent further diversification via a GC response. These findings suggest that the presence of CD27⁺IgD⁺IgM⁺ B cells does not preclude class switching and affinity maturation within the spleen or tonsils in response to *S. pyogenes* infection, a key conceptual advance that demonstrates how protective and durable responses can form even in the presence of pre-existing unswitched memory. Therefore, how polysaccharide vaccines are formulated and delivered may profoundly influence immune response outcome. We posit that conjugation to a protein carrier may lead to different anti-glycan Ab response kinetics and isotype distributions to those of a live or live-attenuated bacterial vaccine, with implications for the quality and durability of immunity generated. In the latter case, our data suggest that vaccination strategies should consider patient age, as children may require multiple doses to develop an adult-like immune repertoire capable of providing effective protection, and this may coincide with the capacity to bypass the marginal zone response to glycans. Thus, our observations outline a molecular blueprint for overcoming non-protective marginal zone responses to generate a diverse, GC-derived, and class-switched Ab response upon vaccination.

Polysaccharide-containing vaccines are generally ineffective at generating durable Ab responses in infants but the underlying mechanisms remain unclear. For example, when the MenC meningococcal vaccine is given to infants under 12 months of age, only a small proportion of vaccinated infants remain seroprotected four years later^56^, while Ab responses in adolescents are more durable^62^. In this study, tonsillar memory and GC B cells in children aged 2–12 years were comparable between glycan- and protein-specific antigen targets, while glycan-specific cells were notably underrepresented among surface BCR-expressing ASCs. This could be due to differences in differentiation, proliferation, or survival between glycan- and protein-specific ASCs although whether these cells arise directly from memory B cells via extrafollicular plasmablast differentiation or through the GC remains unknown. However, the clonal sharing patterns—biased toward GC over memory—and the higher mutation rates observed in ASCs compared to memory B cells in this study suggest that most of the glycan-specific ASCs originate from the GC, so driving this response would potentially be beneficial for early childhood immunity. Mechanistically, glycan-specific GC B cells exhibited lower expression of the IL-21 receptor, a key signal for continued GC participation and ASC differentiation^30,33^. The decline in Ab levels observed in children following Hib glycoconjugate vaccination^3^ suggests that even with a T cell-engaging protein carrier, glycan-specific B cell responses may be insufficient to generate long-lived ASCs. Our findings align with this phenomenon: despite participating in a GC, glycan-specific B cells exhibited reduced expression of antigen presentation machinery, potentially limiting their ability to engage with T cells. Vaccination strategies that enhance T cell support to glycan-specific B cells within the GC may thus be key to improving the durability of polysaccharide vaccine-induced immunity.

Overall, the findings reported here refine the molecular rules that govern B cell proliferation, survival and differentiation in antigen-specific GC responses, highlighting that glycan-specific responses can readily engage this canonical program.

## Resource availability

### Lead contact

Further information and request for resources and reagents should be directed to and will be fulfilled by the lead contact, Danika Hill

### Materials availability

Reagents generated in this study are available from the lead contact with a completed Material

### Data and code availability

Raw and processed single-cell sequencing data are available online at GEO. Code, processed data and sample metadata can be found at <u>10.5281/zenodo.15062289</u>, currently under embargo.

## Supporting information

Supplementary datafile 1

Supplementary datafile 2

## Acknowledgments

We thank all volunteers who were involved in tonsillectomy collections, along with the donors, families and friends of spleen organ donors. We thank all volunteers that participated in the CHIVAS-M75 study, along with study doctors, nurses and support staff. We thank Dr Danilo Gomes Moriel and GSK Vaccines Institute for Global Health for supplying GAC. GSK Vaccines Institute for Global Health is an affiliate of GlaxoSmithKline Biologicals SA. We thank the NSW Health Organ and Tissue Donor Service for providing access to cadaveric spleen tissues, and the Australian Red Cross Lifeblood for providing access to buffy packs. We thank members of the Monash University - Alfred Research Alliance Flow Cytometry Platform (ARAFlowCore) for expert assistance and training with flow cytometry and cell sorting, and the Monash University - School of Translational Medicine (STM) Genomics Platform for assistance with sequencing.

This study was funded by a Michelson Prize from the Human Immunome Project and Michelson Medical Research Foundation (D.L.H.), a National Institute of Health R01 Grant (1R01AI173567-01, A.C.S), Australian National Health and Medical Research Council Ideas Grant (GNT2038630, D.L.H), an Australian Government Research Training Program (RTP) Scholarship (H.A.F.), Australian National Health and Medical Research Council Early Career or Investigator Fellowships (J.O., D.L.H., A.C.S., S.W.S., S.G.T.), the Maurice Wilkins Centre for Molecular Biodiscovery (3716490, N.J.M.), and a Chan Zuckerberg Initiative Single-Cell Biology grant (2021-237883, S.S., M.N.). J.O. is also supported by a Melbourne Children’s Campus Clinician–Scientist fellowship and A.C.S. by a Viertel Senior Medical Research Fellowship. The CHIVAS-M75 study was funded by the Australian National Health and Medical Research Council (1099183). M.J.B. is supported by Snow Medical Foundation Fellowship (2022/SF167) and CSL Centenary Fellowship. The Burnet Institute is supported by the NHMRC for Independent Research Institutes Infrastructure Support Scheme and by Victoria State Government Operational Infrastructure Support.

## Author Contributions

Conceptualization D.L.H., M.J.B., H.R.F.; Data curation D.L.H., H.A.F., C.P., N.K., S.B.; Formal analysis D.L.H., H.A.F., C.P., N.K., S.B., J.N.; Funding acquisition D.L.H., J.O., A.C.S., N.J.M.; Investigation D.L.H., H.A.F., C.P., T.N., A.L.W.; Methodology J.N., N.K., S.B.; Resources M.J.B., A.C.S., H.R.F., N.J.M., K.C., N.C., G.L., B.W.B., N.S., J.B., S.V., K.G., L.G., E.L., K.D., S.G.T., P.S.L., S.W.S., M.N., S.S., J.O.: Supervision D.L.H., H.R.F., J.O., A.C.S., D.M.T.; Visualization D.L.H., H.A.F., C.P., N.K., S.B.; Writing – original draft D.L.H., H.A.F., C.P.; Writing – review & editing H.A.F., C.P., H.R.F., S.B., N.K., P.S.L., T.N., K.C., N.C., A.L.W., J.H., D.A., G.L., B.W.B., I.Q., M.J.R., N.S., J.B., S.V., K.G., L.B., E.L., K.D., S.G.T., J.N., N.J.M., S.W.S., M.N., S.S., J.O., D.M.T., M.J.B., A.C.S., D.L.H.

## Materials and Methods

### Human Ethics

Healthy adult buffy coat samples were obtained from blood donors at the Australian Red Cross Lifeblood, with approval from Monash University Human Research Ethics Committee. Normal human spleens were obtained from organ donors (NSW Organ and Tissue Donation Service) with approval from St Vincent’s Hospital HREC and The Alfred Hospital HREC (284/23). Tonsil tissue was collected following routine tonsillectomy in both Queensland and Melbourne, Australia, with approval through the Queensland Children’s Hospital HREC (HREC/22/QCHQ/81365) for tissue collected through Queensland Children’s Hospital, Royal Brisbane and Women’s Hospitals, and Surgical Treatment and Rehabilitation Service, Queensland, and approval for collection in Melbourne from the Royal Children’s Hospital Melbourne HREC (#88144). Additional local approval for immunology studies was provided by the Alfred Hospital HREC (#607/23 and #284/23). The details of the CHIVAS-M75 study protocol, *emm*75 *S. pyogenes* challenge strain, and trial results have been described previously^41,63,64^. The study, collection of samples, and related immunology research was approved by the Alfred Hospital HREC (#507/17). Previously published data^21^ of serum IgG against SpyCEP and GAC were reanalyzed in this study. Written informed consent was provided by all donors or participants or from their legal guardian/next of kin.

### Sample processing

PBMCs were collected by Ficoll-Hypaque density centrifugation or using CPT tubes. Human spleens were processed to single-cell suspensions by pushing through a cell strainer followed by Ficoll-Hypaque (GEHealthcare, Chicago, IL, USA) separation to isolate the mononuclear cell layer. Tonsil samples were processed after dissection by incubating in a solution of Ham’s F12 nutrient mix, Penicillin-Streptomycin, and either Primocin or Normocin for 1 hr at 4 °C, rinsing in PBS, then mechanically dissociating through an 100 µM strainer in a solution of RPMI (+glutamax or L-glutamine), 10% FBS, MEM non-essential amino acids, sodium pyruvate, Penicillin-Streptomycin, Insulin-Transferrin-Selenium, and Primocin or Normocin. The resulting suspension was frozen in FBS+10% DMSO. To determine *S. pyogenes* colonisation, whole tonsil on ice was incubated in RPMI, then media containing the tonsil was diluted in Todd-Hewitt broth and incubated overnight at 37 °C with 5% CO_2_. Gram positive beta-haemolytic colonies were selected by streaking 20 µL of culture onto Columbia nalidixic acid agar and incubating overnight at 37 °C with 5% CO_2_, then streaking onto a horse blood agar plate overnight and typed as Group A by latex agglutination test.

### Antigens

Chemically extracted GAC (provided by GSK Vaccines Institute for Global Health) was oxidized as previously described^23^, by incubating with 8 mM sodium periodate in borate buffer for 30 min then quenching with 16 mM sodium sulfite. Oxidized GAC was purified by passing through a PD-10 desalting column, exchanging the buffer for 100 mM sodium acetate, then biotinylated by incubating with biotin-PEG4-hydrazide (at a 3.4:1 w/w ratio) in the presence of 80 mM sodium cyanoborohydride for 48 hrs at 30°C. Excess unreacted biotin was removed by 3 rounds of buffer exchange and stored in PBS with 0.02% sodium azide.

The SpyCEP (D151A and S617A) construct with leader sequence and cell anchoring domain removed, and N-terminal periplasm leader sequence, 6xHis-tag, and AviTag sites, was cloned into the pET30 vector. The SpyCEP protein was produced by inducing expression in transformed *E. coli* BL21 (DE3) with isopropyl ß-D-1-thiogalactopyranoside overnight at 18 °C. The periplasmic fraction was extracted by incubating harvested cells with 20% (w/v) sucrose in PBS, followed by 5 mM EDTA and then 20 mM MgCl_2_. His-tagged protein was purified using a 5 mL HisTrap Excel column (Cytiva), eluted with 400 mM imidazole and further purified via size exclusion chromatography using a Superdex 200 Increase 10/300 GL column (Cytiva) equilibrated in PBS. Purified SpyCEP was enzymatically biotinylated according to established protocols ^65^ and free biotin was removed via size exclusion chromatography using a Superdex 200 Increase 10/300 GL column, equilibrated in PBS. SpyCEP stocks were stored at -80C in PBS+10% (v/v) glycerol.

Influenza A/Michigan/2015 haemagglutinin (HA) protein with Y89F mutation to prevent sialic acid binding, N-terminal leader sequence and a C-terminal biotin ligase AviTag and 6xHis-tag was produced as previously described^66^. Briefly, the HA construct was cloned into a pCR3 plasmid, expressed using the Expi293F Expression System (Thermo Fisher Scientific) and His-tagged protein purified using a cobalt TALON Metal Affinity Resin affinity purification column (Takara Bio) and eluted with imidazole. Purified protein was biotinylated using the BirA500 biotin-protein ligase kit (Avidity) and dialysed against 10 mM Tris.

### Tetramerization and staining concentrations

Biotinylated GAC, SpyCEP, and HA were tetramerized by the addition of fluorochrome-conjugated streptavidin at a 1:4 streptavidin:protein molar ratio. A tenth of the required volume of streptavidin was added every 5 min for a total of 10 additions. For each Ag, tetramers were made using two different fluorochromes to allow for dual-discrimination gating to reduce the inclusion of non-specific binding cells. Cells were stained with Ag tetramers at 1.25 pmol of tetramer per 10x10^6^ cells in a final staining volume of 200 µL.

### Flow cytometry and cell sorting

Frozen mononuclear PBMCs, tonsil or spleen cells were thawed at 37°C in thawing media (RPMI + 10% FCS + 1% Pen-Strep), then washed and resuspended in flow buffer (PBS + 2% FCS + 2 mM EDTA) and stained with fluorescently labeled Abs (including CD3, CD19, CD20, CD21, CD27, CD95, IgD, IgG, IgM and IgA), viability dye, and Ag tetramers with 1 mM excess biotin. Cells were fixed with BD fixation buffer, filtered through a 70 µm mesh, and analyzed on a Cytek Aurora flow cytometer (Cytek Biosciences).

Samples used for sorting Ag-specific B cells for downstream single-cell sequencing were thawed as above and stained with Ag tetramers on PE and a second fluorochrome for each Ag. Tetramer-binding cells were magnetically enriched using an EasySep PE Positive Selection Kit II (StemCell Technologies). Ag tetramer-binding cells were then stained with fluorescently labeled Abs (CD3, CD19, CD20, CD27, IgD), viability dye, and an anti-human cell surface hashtag Ab (TotalSeq-C, BioLegend) to allow later identification of Ag specificity. Ag tetramer-binding isotype-switched B cells were sorted on a Cytek Aurora CS (Cytek Biosciences). These cells were gated as CD19+ CD3– IgD– viable single lymphocytes positive for both fluorochromes of an Ag tetramer pair, to reduce the inclusion of non-specific binding cells.

### Single cell sequencing

Sorted Ag tetramer-binding isotype-switched B cells were pooled, spun down, resuspended in PBS + 2% FCS, and counted. 40,000 cells were loaded in two separate reactions on a 10X Chromium Next Gem Chip K. VDJ, Gene Expression, and Feature Barcode libraries were prepared using the Chromium Next GEM Single Cell 5’ v2 dual index kit according to the manufacturer’s instructions and sequenced on a NovaSeq 6000 using an S1 kit.

### Bioinformatics

Raw sequence read quality was assessed using FastQC software. Data was demultiplexed using VireoSNP including: cellular genotyping with cellSNP-lite v1.2.3 (Huang et al.) and 7.4 SNPs (with MAF >0.05) from the 1000 genomes project; donor deconvolution with Vireo v0.5.6 (Huang et al). Data were then aligned with CellRanger multi (version 7.2.0) which allows for simultaneous profiling of the V(D)J repertoire, cell surface protein, Ag, and gene expression (GEX) data. The reads were mapped to the Human GRCh38 (GENCODE v32/Ensembl98) reference genome, and the VDJ sequences were mapped to vdj-GRCh38-alts-ensembl-5.0.0 reference transcriptome.

### Data Processing and Analysis

The counts generated were processed and analyzed using the Seurat package (version 5.1.0) in R (version 4.4.0). Counts were filtered to remove low gene expression signals, low-quality cells, and mitochondrial content. Data normalization and scaling were performed using Seurat’s NormalizeData and ScaleData functions. Highly variable genes were identified using the FindVariableFeatures function with the “vst” method. Integration of samples was performed using Seurat’s FindIntegrationAnchors and IntegrateData functions. Dimensionality reduction was performed using Seurat’s RunPCA and RunUMAP functions. Cells were clustered based on gene expression profiles using FindNeighbors and FindClusters functions, with a resolution parameter of 1.8 resulting in 17 clusters. FindIntegrationAnchors was used with the HCATonsil Atlas^36^ to annotate clusters and identified 16,109 anchors from the HCATonsil GCBC dataset, 13,432 anchors and HCAT NMBC dataset. MapQuery was then used with reference.reduction set as “pca” and reduction.model set as “umap” to infer cluster annotations. Marker gene analysis between clusters was conducted using the FindMarkers and FindAllMarkers functions, employing the “roc” test to annotate unidentified cell clusters in the data. The aggregated counts generated by AggregateExpression function were used for pseudo-bulk differential expression analysis, comparing cell types between GAC and SpyCEP conditions. The differential testing was performed using edgeR v4.4.1 and limma v3.62.2. Visualization of the data was performed using ggplot2 (version 3.5.1) and Seurat’s plotting functions. Data from CellRanger (e.g. contig, feature) was further processed through dandelion v0.5.0 for BCR analysis (Suo et al.), which is used to study immune repertoires, clonal expansion and B cell diversity. Briefly, data were prepared using the singularity preprocessing step (V(D)J reannotation with igblastn, reassigning heavy chain IG V gene alleles with TlgGER, reassigning IG constant region calls, quantifying mutations with SHazaM), gene expression and VDJ data processed via Scanpy and V(D)J analysis performed with Dandelion to define chain status, clone size, VDJ/isotype status usage etc. scRepertoire v2.2.1 was used to visualise clonal expansion and B cell diversity.

### Statistical analysis

*P* values were determined using two-sided non-parametric methods; including Wilcoxon signed-rank test for paired two-group comparisons, Dunn’s test for 3+ group comparisons. Where repeated two group comparisons were performed, *p* values were adjusted for multiple corrections using the false discovery rate approach. Correlation coefficients and correlation *p* values were determined using Spearman’s method (two-sided) with a linear regression line included to indicate linear trends.

