## Supplementary material for "Antibody responses against bacterial glycans affinity mature and diversify in germinal centers"

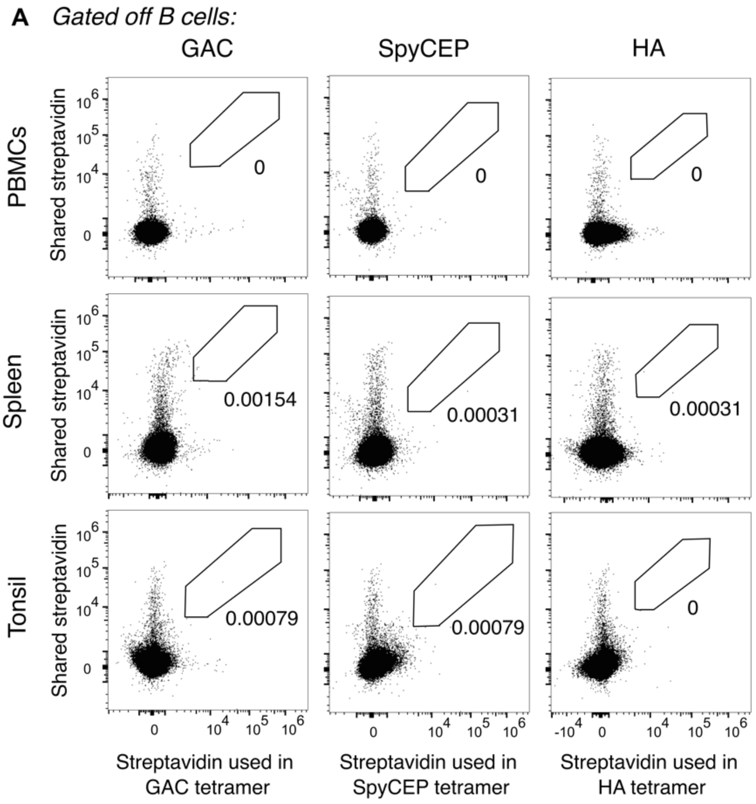


**Supplementary Fig. 1: Streptavidin-only control staining.** (**A**) Representative plots of adult PBMCs, adult spleen, and pediatric tonsil stained with free fluorescent streptavidins (not conjugated to antigen) matching those used in GAC, SpyCEP, and HA tetramers, showing background non-antigen-specific binding of B cells.

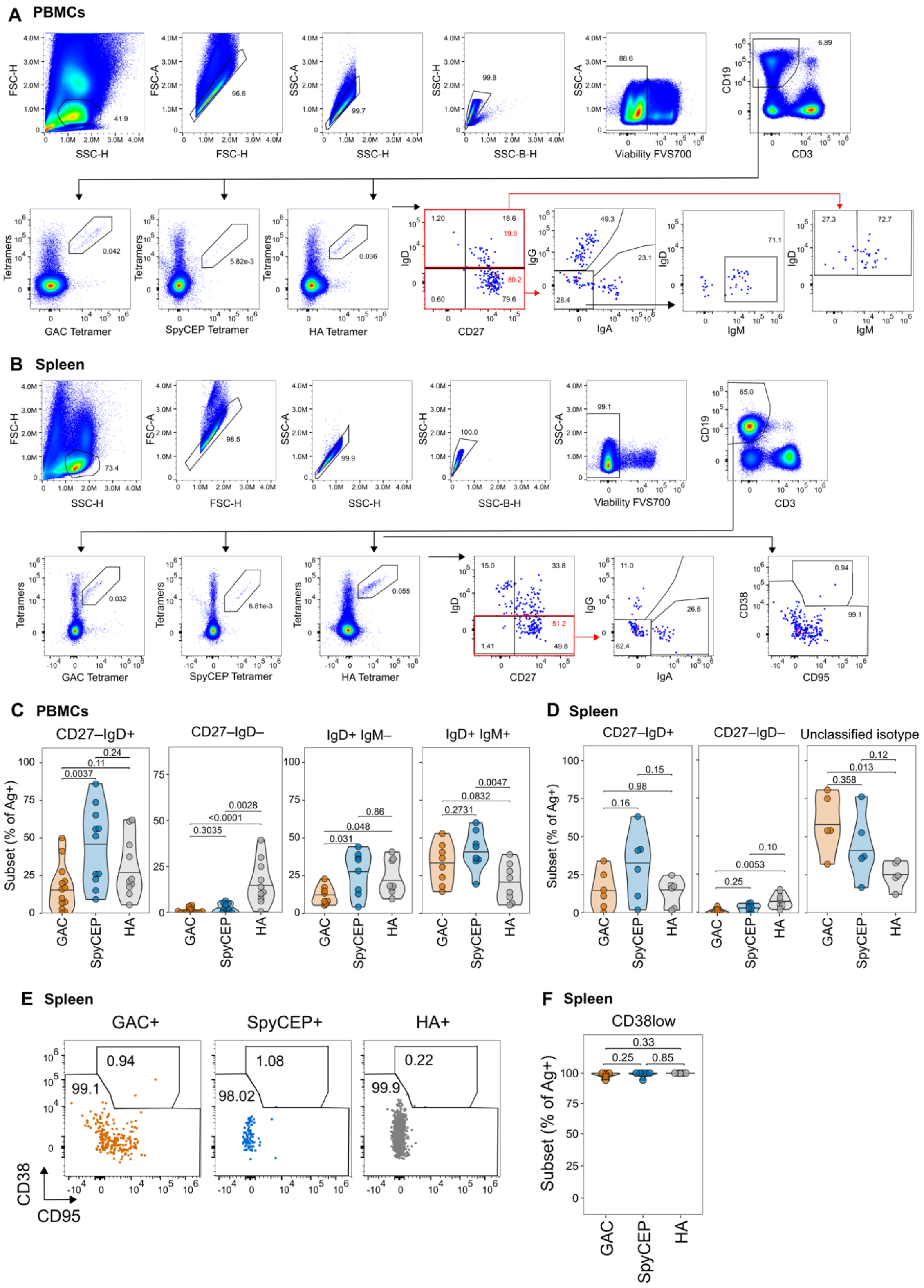


**Supplementary Fig. 2: Gating strategies and additional data for flow cytometric analysis of adult PBMCs and spleens.** (**A-B**) Gating strategy on PBMCs (**A**) or spleens (**B**). Cells were gated as lymphocytes, singlets, live cells, then B cells gated as CD19+CD3-. Antigen specific cells were gated off B cells, then within each antigenbinding population memory and naïve subsets (CD27 and IgD), isotypes, and GC phenotype (CD38 and CD95, spleen only) were gated. Representative gating off GAC+ cells off one sample per panel. (**C**) Frequencies of CD27–IgD+, CD27–IgD–, IgD+IgM–, and IgM+IgD+ cells within GAC+, SpyCEP+, or HA+ B cells in adult PBMCs. (**D**) Frequencies of CD27–IgD+, CD27–IgD–, and unclassified isotype (not IgD+, IgA+, or IgG+) cells within GAC+, SpyCEP+, or HA+ B cells in adult spleens. Horizontal bars in violin plots indicate median values. *P* values were determined using two-sided non-parametric Dunn's test with false discovery rate p-value adjustment for multiple comparisons across each population.

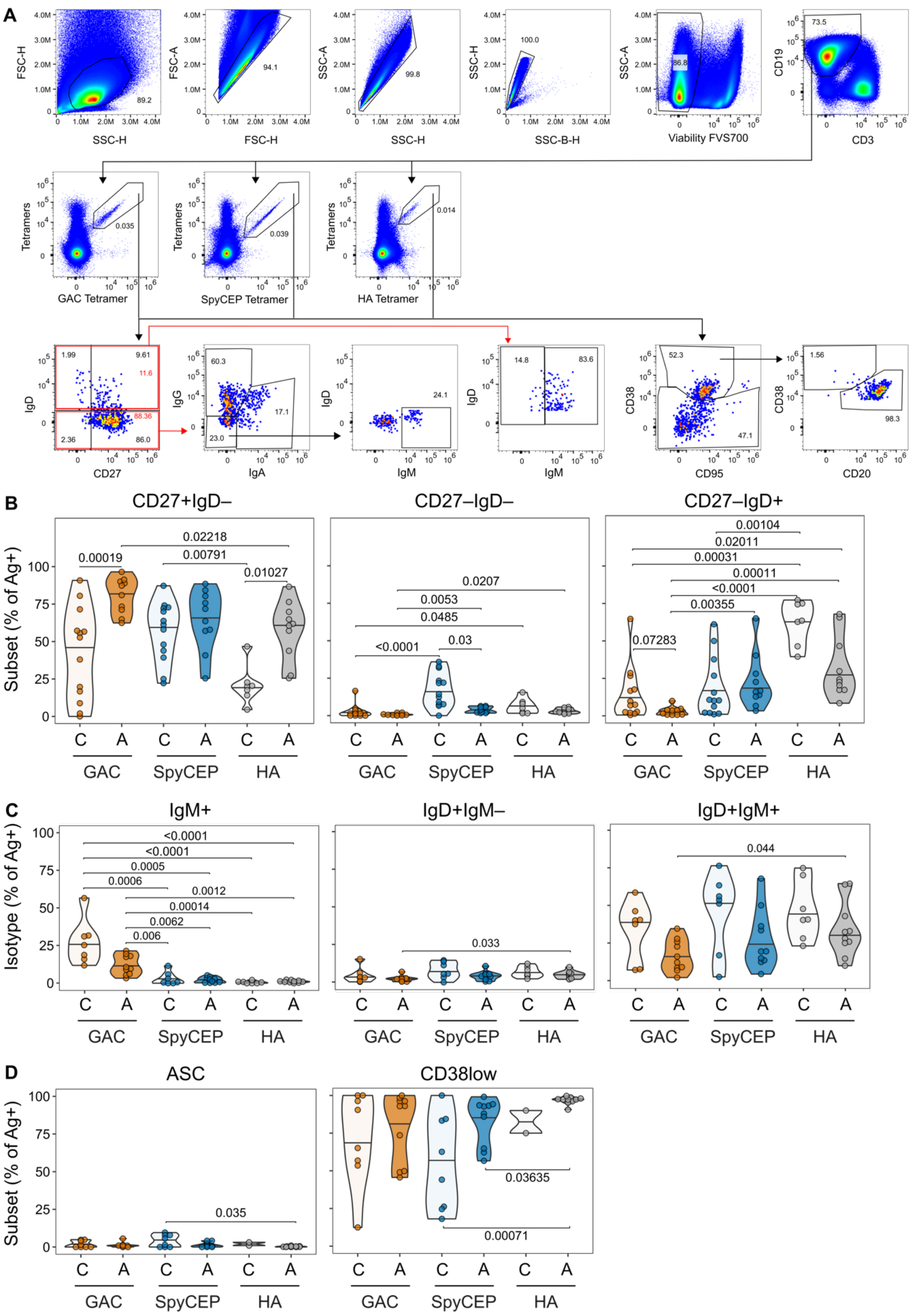


**Supplementary Fig. 3: Gating strategies and additional data for flow cytometric analysis of adult and child tonsils.** (**A**) Gating strategy. Cells were gated as lymphocytes, singlets, live cells, then B cells gated as CD19+CD3-. Antigen specific cells were gated off B cells, then within each antigen binding population cells gated for memory and naïve subsets (CD27 and IgD), isotype (either IgD vs IgM then IgG vs IgA or IgD- then IgG vs IgA), and GC phenotype (CD38 and CD95 for CD38low cells, and CD38+ cells gated as GC phenotype (CD38+ CD20+) or ASC phenotype (CD38++ CD20low). Representative gating off GAC+ cells off one colonised juvenile tonsil sample (SpyCEP and HA not shown). (**B**) Frequencies of CD27+IgD-, CD27-IgD+, and CD27-IgD- cells within GAC+, SpyCEP+, or HA+ B cells for children and adults. Only p <0.05 shown, and only comparisons between antigens or child and adult. Frequencies of CD27+IgD- cells shown in Figure 3J. (**C**) Frequencies of IgM+, IgD+, and IgM+IgD+ cells within GAC+, SpyCEP+, or HA+ B cells for children and adults. Frequencies of IgG+, IgA+ cells shown in Figure 3I. (**D**) Frequencies of ASC (CD38++, CD20low) and CD38low cells within GAC+, SpyCEP+, or HA+ B cells. Frequency of GC B cells shown in Figure 3I. Horizontal bars in violin plots indicate median values. *P* values were determined using two-sided non-parametric Dunn’s test with false discovery rate p-value adjustment 15 pairwise comparisons. Only p <0.05 shown.

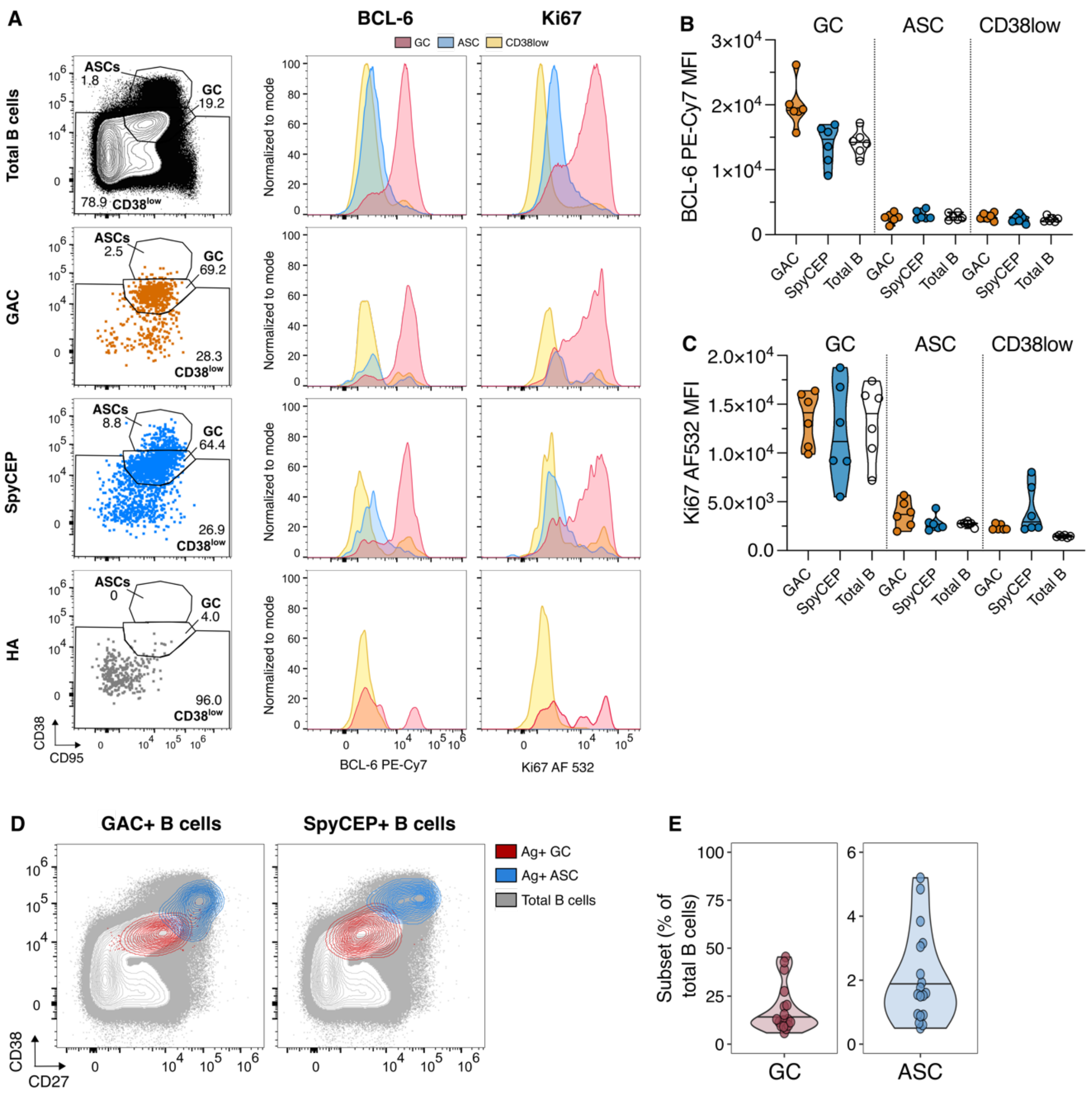


**Supplementary Fig. 4: Confirmation of GC B cell phenotypes.** (**A**) Intranuclear staining of BCL-6 and Ki67 in total and Ag+ GC, ASC, and CD38low cells. (**B-C**) MFI of BCL-6 (**B**) and Ki67 (**C**) staining on total and Ag+ B cell subsets. (**D**) Representative plot showing the expression of CD38 and CD27 by GAC+ or SpyCEP+ GC B cells and ASCs. compared to total B cells. (**E**) Frequencies of GC B cells and ASCs within total B cells in pediatric tonsils. Horizontal bars in violin plots indicate median values.

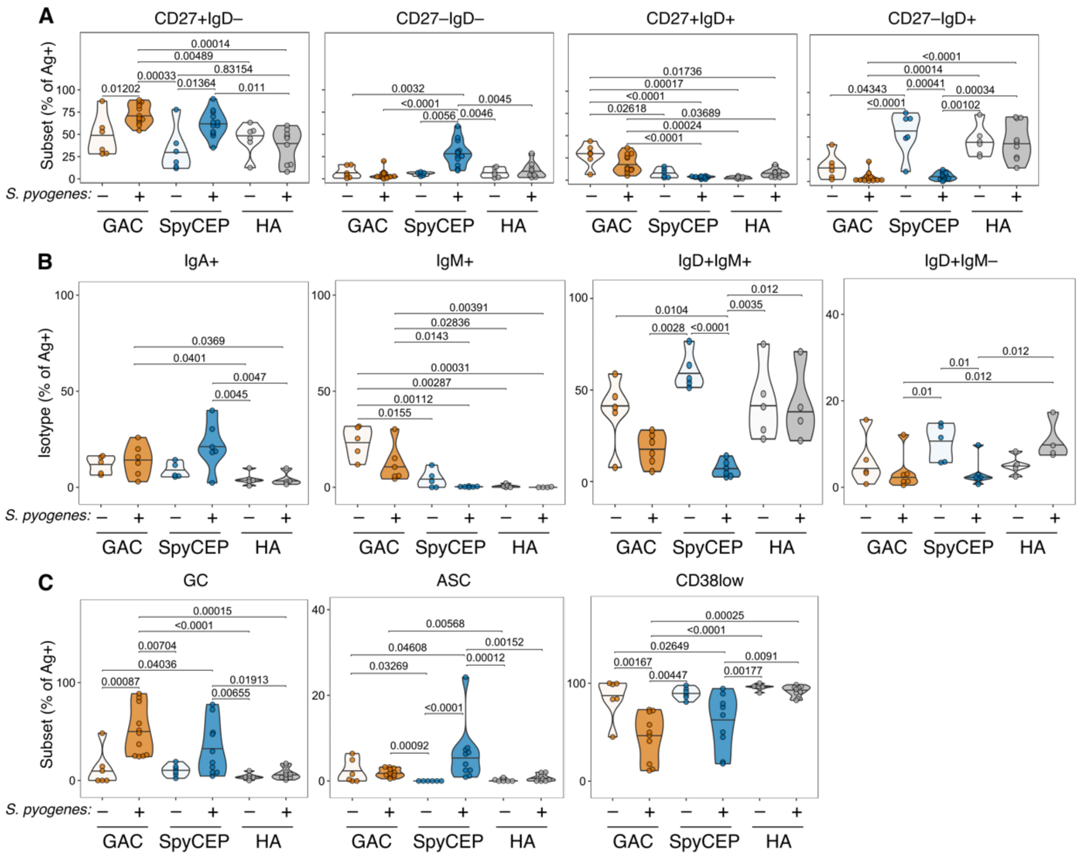

**Supplementary Fig. 5: Additional data from flow cytometry analysis of tonsils with and without *S. pyogenes* colonization.** (**A**) Frequencies of CD27+IgD+, CD27+IgD-, CD27-IgD+, and CD27-IgD- cells within GAC+, SpyCEP+, or HA+ B cells for tonsils with and without *S. pyogenes* colonisation. (**B**) Frequencies of IgA+, IgM+, IgD+, and IgM+IgD+ cells within GAC+, SpyCEP+, or HA+ B cells for tonsils with and without *S. pyogenes* colonisation. (**C**) Frequencies of GC (CD38+, CD20+), ASC (CD38++, CD20low) and CD38low cells within GAC+, SpyCEP+, or HA+ B cells. Horizontal bars in violin plots indicate median values. *P* values were determined using two-sided non-parametric Dunn's test with false discovery rate p-value adjustment for 15 pairwise comparisons. Only p <0.05 shown.

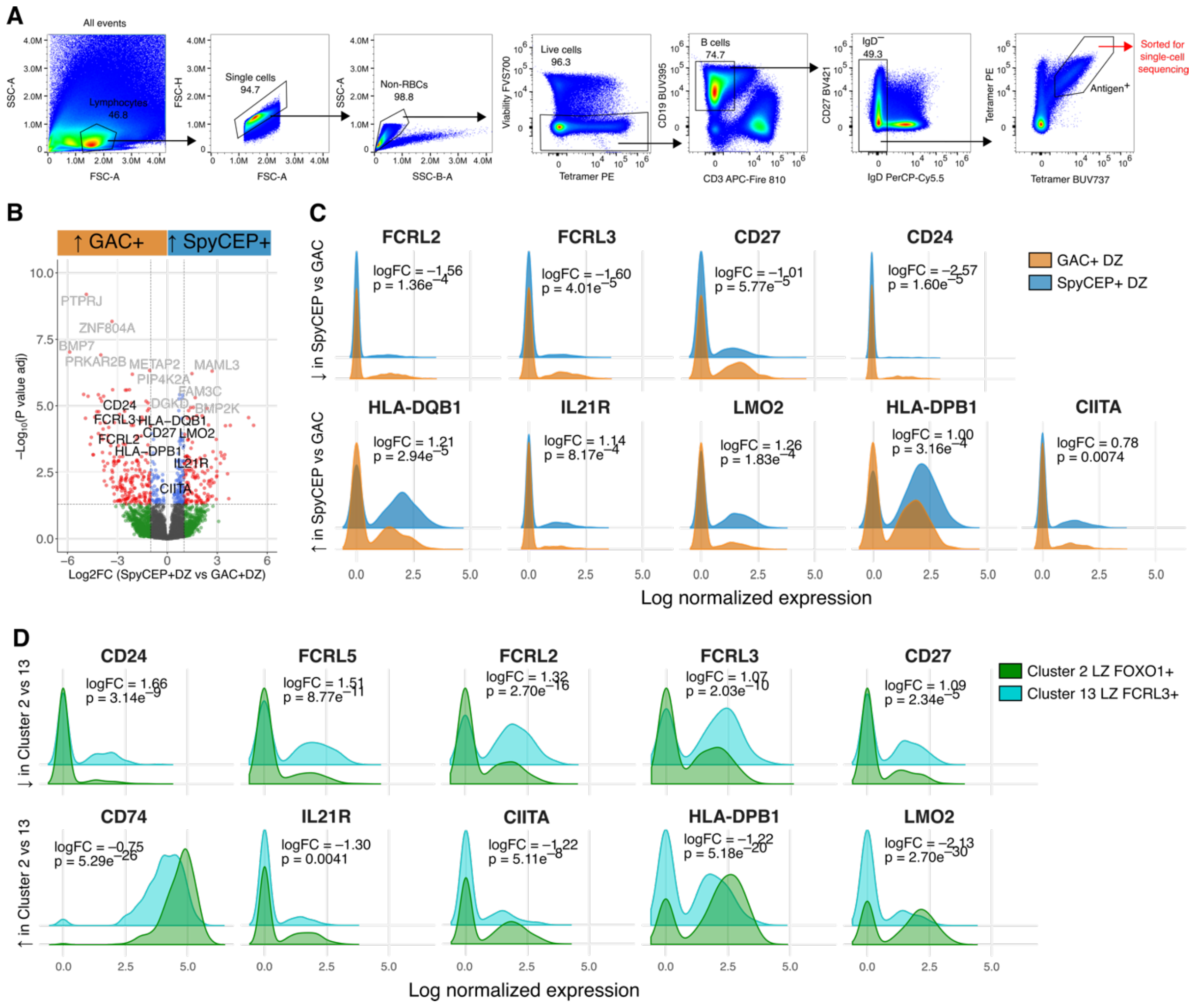


**Supplementary Fig. 6: Single-cell sequencing B cell sort strategy and additional data.** (**A**) Gating strategy for sorting Ag+ IgD– B cells from pediatric tonsils with *S. pyogenes* colonization. (**B**) Volcano plot showing differentially expressed genes between SpyCEP+ DZ B cells and GAC+ DZ B cells. (**E-F**) Highlighted differentially expressed genes between GAC+ and SpyCEP+ DZ B cells (**E**), and Cluster 2 (LZ FOXO1+) and Cluster 13 (LZ FCRL3+) cells (combined GAC+ and SpyCEP+ cells) (**F**).

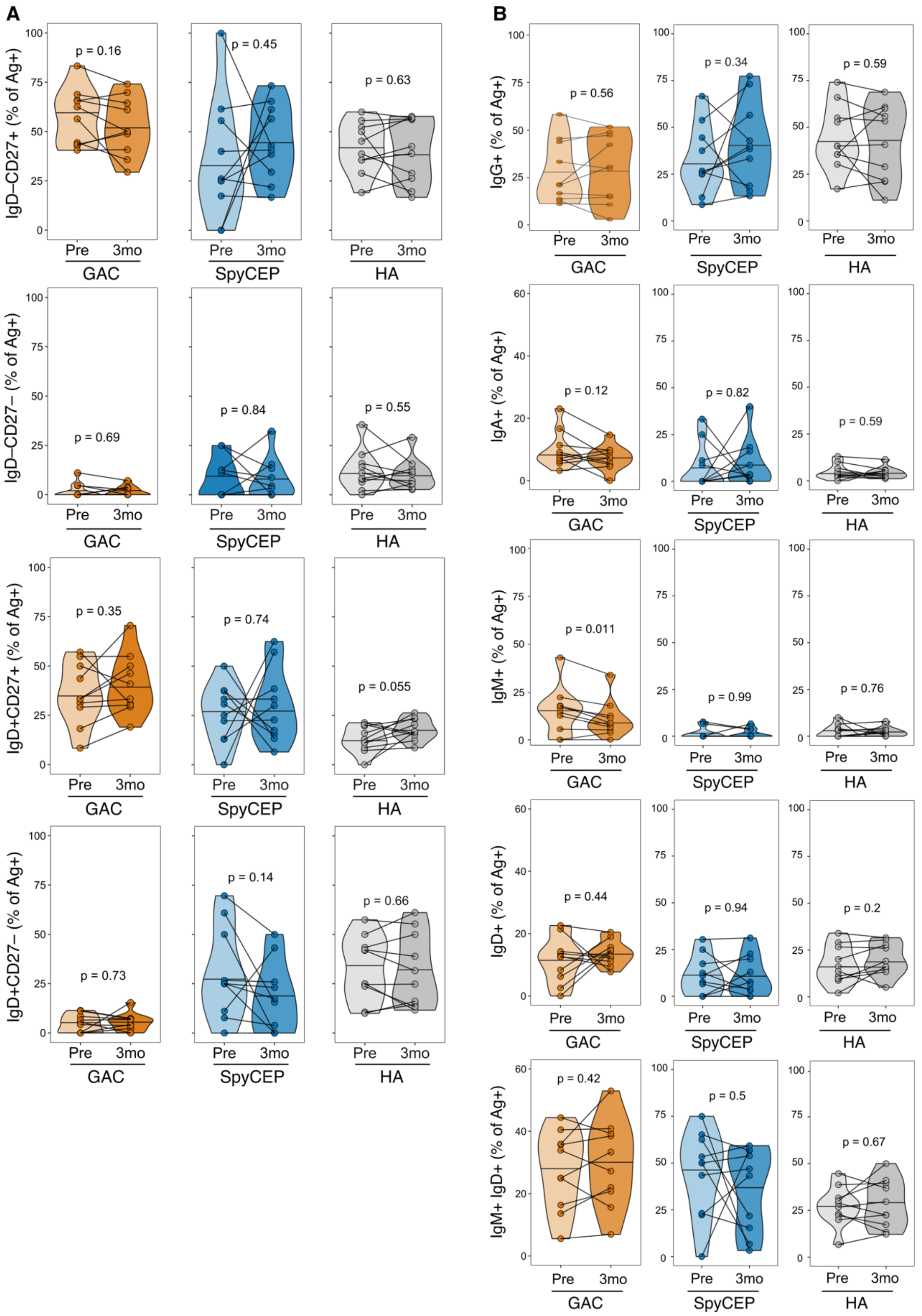


**Supplementary Fig. 7: Additional data from flow cytometric analysis of *S. pyogenes* human challenge PBMCs.** (**A**) Frequencies of CD27+IgD+, CD27+IgD-, CD27-IgD+, and CD27-IgD- cells within GAC+, SpyCEP+, or HA+ B cells from PBMC samples pre- and 3mo post- *S. pyogenes* challenge. (**B**) Frequencies of IgG+, IgA+,

IgM+, IgD+, and IgM+IgD+ cells within GAC+, SpyCEP+, or HA+ B cells pre- and 3mo post-challenge. Horizontal bars in violin plots indicate median values. *P* values were determined using two-sided Wilcoxon signed-rank test for paired two-group comparisons.
